# Loss of tristetraprolin activates NF-κB induced phenotypic plasticity and primes transition to lethal prostate cancer

**DOI:** 10.1101/2022.08.05.500896

**Authors:** Katherine L. Morel, Anis A. Hamid, Beatriz G. Falcón, Jagpreet S. Nanda, Simon Linder, Andries M. Bergman, Henk van der Poel, Ingrid Hofland, Elise M. Bekers, Shana Trostel, Scott Wilkinson, Anson T. Ku, Deborah L. Burkhart, Minhyung Kim, Jina Kim, Jasmine T. Plummer, Sungyong You, Adam G. Sowalsky, Wilbert Zwart, Christopher J. Sweeney, Leigh Ellis

**Affiliations:** South Australian Immunogenomics Cancer Institute, University of Adelaide, Adelaide, SA, Australia; Department of Medical Oncology, Dana-Farber Cancer Institute, Harvard Medical School, Boston, MA, USA; Department of Surgery, University of Melbourne, Melbourne, Australia; Division of Hematology and Oncology, Department of Medicine, Cedars-Sinai Medical Center, Los Angeles, CA; Division of Oncogenomics, Oncode Institute, The Netherlands Cancer Institute, Amsterdam, the Netherlands; Division of Medical Oncology, The Netherlands Cancer Institute, Amsterdam, The Netherlands; Division of Urology, The Netherlands Cancer Institute, Amsterdam, The Netherlands; Core Facility Molecular Pathology and Biobanking, The Netherlands Cancer Institute, Amsterdam, The Netherlands; Division of Pathology, The Netherlands Cancer Institute, Amsterdam, The Netherlands; Laboratory of Genitourinary Cancer Pathogenesis, Center for Cancer Research, National Cancer Institute, National Institutes of Health, Bethesda, MD, USA; Department of Biomedical Sciences, Cedars-Sinai Medical Center, Los Angeles, CA, USA; Department of Developmental Neurobiology, St. Jude Children’s Research Hospital, Memphis, TN, USA; Division of Urology, Department of Surgery, Cedars-Sinai Medical Center, Los Angeles, CA, USA; Cedars-Sinai Center for Bioinformatics and Functional Genomics, Los Angeles, CA; Cedars-Sinai Samuel Oschin Comprehensive Cancer Institute, Los Angeles, CA

## Abstract

Phenotypic plasticity is a hallmark of cancer and increasingly realized as a mechanism of resistance in androgen indifferent prostate tumors. It is critical to identify mechanisms and actionable targets driving phenotypic plasticity. Here, we report that loss of tristetraprolin (TTP, gene *ZFP36*), an RNA binding protein that regulates mRNA stability increases NF-κB activation and is associated with higher rates of aggressive disease and early recurrence in primary prostate cancer (PCa). We examined the clinical and biological impact of *ZFP36* loss combined with *PTEN* loss, a known driver of PCa. Combined loss of *PTEN* and *ZFP36* expression was associated with increased risk of recurrence in multiple independent primary PCa cohorts, and significantly reduced overall survival and time to progression following castration in genetically engineered mouse models. *ZFP36* loss alters the cell state that is driven by *PTEN* loss, demonstrated by positive enrichment of gene sets including EMT, inflammation, TNFα/NF-κB, IL6-JAK/STAT3. *ZFP36* loss also induces enrichment of multiple gene sets involved in cell migration, chemotaxis, and proliferation. Use of the NF-κB inhibitor dimethylaminoparthenolide induced significant therapeutic responses in tumors with *PTEN* and *ZFP36* co-loss and reversed castration resistance. This work identifies a novel molecular mechanism driving phenotypic plasticity and castration resistance through loss of *ZFP36* expression, that can be reversed by inhibition of NF-κB activity.

## Introduction

Multiple factors have been shown to drive prostate cancer (PCa) progression including inflammation (1, 2) and NF-κB (p65) activation (3, 4). Chronic inflammation is commonly observed in prostate tumors, although the exact stimuli required to initiate and maintain prostatic inflammation are not fully understood. This inflammatory response consists of the recruitment and expansion of leukocytes including myeloid cells, macrophages and lymphocytes in the prostate (5–7). Immunohistochemical analysis has shown p65 is active in early PCa, including prostate intraepithelial neoplasia (PIN) and low- and high-grade PCa (8, 9). *In vitro* and *in vivo* modeling have linked constitutive NF-κB activation to many of the hallmarks of cancer including proliferation and evasion of apoptosis which can be abrogated by NF-κB inhibition (10, 11).

Due to the complex feedback control mechanisms, markers of NF-κB activation that associate with poor PCa outcomes have been elusive (12). With a systems biology approach focused on identifying regulators associated with development of lethal PCa, we described tristetraprolin (TTP, gene name *ZFP36*), as a key node in controlling NF-κB activation and progression to lethal PCa (13). We noted that patients with lower *ZFP36* were more likely to relapse and die of PCa (14–17) and that silencing *ZFP36* made non-transformed prostate cells (RWPE-1) proliferate more rapidly, while overexpressing *ZFP36* decreased proliferation of the LNCaP PCa cell line (14).

Tristetraprolin, like butyrate response factors, is a member of the TPA-inducible sequence 11 (TIS11) family of RNA-binding proteins that directly binds to *cis*-acting adenosine and uridine-rich elements (AREs) in the 3′ UTR of target mRNA leading to recruitment of enzymes for the rapid shortening of the poly(A) tail which results in transcript de-adenylation and degradation (18, 19). This posttranscriptional regulation of mRNA stability allows cells to respond to intracellular and extracellular stimuli. Relevant to our work is that TTP also mediates degradation of TNF-α mRNA (an NF-κB activator) and IL-1β (20). Loss of TTP therefore results in NF-κB hyper-activation with associated inflammatory conditions including arthritis and dermatitis (21–23). Inflammation plays a crucial role during PCa initiation, progression, and metastasis (24). Chronic elevation of pro-inflammatory genes is known to promote tumor evolution by increased proliferation, angiogenesis, metastasis, survival, and drug resistance of cancer cells (25, 26). Because TTP can negatively regulate many inflammatory and oncogenic cytokines (27), this inflammatory suppressive feature of TTP may prevent the occurrence and progression of many cancers including PCa. In addition to mediating expression of inflammatory genes, TTP has also been demonstrated to mediate degradation and control expression of oncogenes and tumor suppressor genes including *c-Myc*, *c-JUN* and *p53* (19). Moreover, neuronal differentiation exhibited dependence on TTP expression and function (19, 28), implying that TTP may be an important player in determining cell identity and tumor evolution.

TTP has previously been described as a ‘prospective’ cancer tumor suppressor (18, 29, 30).Here, we establish the role of TTP as a tumor suppressor in PCa and determine NF-κB activation contributes to the deleterious effects of TTP loss within tumor cells. Given the finding that TTP loss occurs early in PCa development, we used genetically engineered mouse models (GEMMs) to investigate the impact of TTP loss in combination with PTEN loss on prostate epithelial cell transformation to cancer. Mice with co-loss of TTP and PTEN had a significant increase in PCa progression and a decrease in responsiveness to castration compared to GEMMs with PTEN loss alone. We demonstrate that this aggressive phenotype, driven by TTP loss, induces phenotypic plasticity, and that this altered plasticity occurs independent of RB1 and P53. This is concluded by the observed decreased androgen receptor expression and function and de-repression of gene signatures enriched for nervous system development and leukocyte cell identity and function. Our data provides new insight towards mechanistic understanding of phenotypic plasticity involving a TTP-p65 signaling axis that can be countered by pharmaceutical targeting of NF-κB. Finally, these data provide strong rationale for the clinical assessment of enhancing the efficacy of hormonal therapy by NF-κB inhibition.

## Methods

### Identification of *ZFP36* gene signature

We defined a *ZFP36* gene signature by identifying differentially expressed genes (DEGs) between two groups. To this end, we first selected samples with high (upper 75%) and low (lower 25%) expression of *ZFP36* in TCGA PRAD data (31). We then identified DEGs between *ZFP36* high and low groups using an integrated hypothesis testing method as previously reported (32). Briefly, 1) T-statistics, rank-sum statistics, and log2-median-ratio between the groups were computed for each gene. 2) Empirical distributions of the null hypothesis were estimated by calculating the three statistics for the genes after randomly permuting the samples 10,000 times. 3) For each gene, three P-values of the observed statistics were computed using their corresponding empirical distributions of null hypothesis by two-tailed test. 4) The three P-values were combined into an overall P-value using Stouffer’s method (33). 5) Lastly, we performed multiple testing correction through Storey’s method (34). DEGs were defined as the genes with an FDR less than 0.05 and absolute log2-median-ratio greater than 0.58 (1.5-fold). The Z-score method was used to quantify signature activation (51 upregulated genes in *ZFP36*-low group) and applied to localized PCa validation cohorts (TCGA PRAD (31); Gulzar *et. al.* (35); Taylor *et. al.* (36); dichotomized by median Z-score) with available clinical outcomes data (biochemical recurrence or disease-free survival), using the Kaplan-Meier method and p-values were calculated by the log-rank test. The HR, upper and lower bound of 95% CI of previously published HSPC cohorts associating TTP expression (by RNA profiling or IHC) with biochemical recurrence (37) were employed for meta-analysis using the *meta* function in STATA version 17.0 (StataCorp, College Station, TX). Meta-analysis was also performed of two previously published cohorts (Health Professionals Follow-up Study (HPFS) and Physicians’ Health Study (PHS) (14); GenomeDx Decipher (38)) correlating *ZFP36* loss with lethal PCa (PCa death or metastases).

### Fluorescent immunohistochemistry and spectral imaging in human tissue microarrays

A retrospective cohort of 112 patients with localized PCa who underwent radical prostatectomy (RP) (39, 40) was annotated for clinicopathologic features, disease outcomes and follow-up. Archival formalin-fixed, paraffin-embedded RP specimens were used to construct three tissue microarrays (TMAs). Fluorescent immunohistochemistry (FIHC) for PTEN was performed using a multiplexed tyramide signal amplification method, with AMACR masking tumor epithelium and a DAPI counterstain. Staining, imaging, quantification, and summarization of PTEN expression was performed as previously described (39), with loss defined as the lower cohort quartile of combined cytoplasmic and nuclear expression. FIHC for TTP was performed for 1 hr employing the same TMAs. The distribution of cytoplasmic TTP expression intensity was quantified at an individual cell level across all cores. Cases with a microarray core exhibiting lower-quartile TTP expression intensity in greater than two-thirds of the tumor cells was deemed to have TTP loss. FIHC antibody details are outlined in Supplementary Table 2.

To examine the clinical effect of compound PTEN and TTP loss, the lower quartile of normalised RNA expression of PTEN and TTP in TCGA PRAD (31) and Taylor et al (36) cohorts defined loss of expression. For these cohorts and DFCI FIHC, the PSA based end-point of disease-free survival was estimated using the Kaplan Meier method and Cox proportional hazards model estimated the hazard ratio (HR) and 95% confidence interval (95% CI); p-values for association tests between gene expression and endpoints were calculated using the log-rank test. In the DFCI FIHC cohort, multivariable analysis was performed adjusting from Gleason score, tumor stage (T stage) and PSA level at diagnosis.

### DARANA clinical data

Patients with localized PCa were enrolled in the DARANA study (Dynamics of Androgen Receptor Genomics and Transcriptomics After Neoadjuvant Androgen Ablation; ClinicalTrials.gov number, NCT03297385) prior to radical prostatectomy. Patients were treated with neoadjuvant enzalutamide for three months prior to prostatectomy, with tumor biopsies taken pre- and post-treatment. Gene expression (RNA-seq) and ChIP-Seq tumor data was generated as described in (41). For IHC analysis, the ENZ-treated patient cohort (n = 50) was matched in a 1:2 ratio to untreated control patients (not receiving ENZ prior to prostatectomy; n = 109) based on clinicopathological parameters (initial PSA, Gleason score, TNM stage, age) using the R package MatchIt (v.4.1.0) (42). Tissue microarrays (TMAs) were prepared containing 3 cores per FFPE tumor sample. Tumor-dense areas in FFPE megablocks were marked by an expert pathologist on a H&E slide. TMAs were stained with antibody for 60 minutes at room. Bound antibody was detected using the OptiView DAB Detection Kit (Ventana Medical Systems) and slides were counterstained with hematoxylin. Antibody details are listed in Supplementary Table 2. TTP staining intensity (negative, low, high) in tumor cells was scored by an expert pathologist.

### NCI clinical data

Patients with intermediate-to-high risk PCa were enrolled in a clinical trial of six months of neoadjuvant ADT plus enzalutamide (43, 44) prior to radical prostatectomy. Usable RNA was obtained from paired pre-and post-treatment tissues (including complete pathologic responders) from 36 patients, using the biopsy (pre-treatment) or whole mount prostatectomy (post-treatment) block best corresponding to the index lesion visualized by multiparametric MRI. Usable baseline biopsy RNA was available from 37 patients. RNA was extracted from ribbon curls without microdissection using the Qiagen FFPE RNeasy kit. Paired-end RNA-seq libraries were generated using the Illumina TruSeq Stranded Total RNA kit with Ribo-Zero/Globin depletion. Raw RNA-seq reads were deposited to dbGaP (phs001938.v3.p1) and processed data were deposited to GEO (GSE183100).

### Animal ethics statement

All animal breeding and experiments were approved by and performed in accordance with the guidelines of the Dana-Farber Cancer Institute Institutional Animal Care and Use Committee (Animal protocol #19-005).

### Generation of GEMM models

Prostate-specific-Pten/*Zfp36* KO mice were achieved by crossing the loxP-flanked Pten mice (Pten^f/f^)(45) with loxP-flanked *Zfp36* mice (*Zfp36*^f/f^)(46) and mice expressing Cre recombinase under the control of the rat probasin gene promoter, which is specific for cells of the epithelial cells of the prostate (PB-Cre4)(47). Homozygous Pten^f/f^ mice and PB-Cre4 mice on a C57BL/6 background were purchased from Jackson Laboratory (Bar Harbor, Maine) (48), and homozygous *Zfp36*^f/f^ mice were a gift from Dr. Perry Blackshear. Heterozygous matings, male PB-Cre4/Pten^+/f^/*Zfp36*^+/f^ crossed with female Pten^+/f^/*Zfp36*^+/f^ mice, were employed to generate prostate-specific Pten/*Zfp36* KO mice and their littermate wild-type (WT) control mice. All experimental mice had Pten homozygous deletion, while *Zfp36* status was examined in WT (PB-Cre4/Pten^f/f^/*Zfp36*^+/+^), heterozygous (PB-Cre4/Pten^f/f^/*Zfp36*^+/f^) or homozygous (PB-Cre4/Pten^f/f^/*Zfp36*^f/f^) states Littermate male mice carrying floxed Pten and *Zfp36* alleles but lacking the PB-Cre4 transgene were used as WT controls.

For PCR genotyping, genomic DNA from ear-notch clips was extracted by alkaline lysis by incubating the tissue sample in 75ul 25mM NaOH/ 0.2 mM EDTA at 95°C for 30 mins, cooling and mixing with 75ul of 40 mM Tris HCl (pH 5.5) to neutralize the reaction. All PCR reactions were run using 2 µL of extracted DNA in GoTaq® PCR Master Mix (Promega) and the following cycling protocol: 95°C for 2 min; 35 cycles of: 95°C for 30 sec, 60°C for 60 sec, and 72°C for 90 sec; 72°C for 5 min. PCR products were visualised on a 1.8% agarose gel (in 1× TAE buffer) containing 1× SYBR Safe DNA dye.

The PB-Cre transgene was detected using the primer pair PbF1 and Cre4R2 to generate a transgene-positive (331 bp) amplicon. Pten-floxed alleles were detected using the primer pair PtenF and PtenR to generate WT (124 bp) or floxed (321 bp) amplicons. *Zfp36*-floxed alleles were detected using the primer pair Zfp36F1 and Zfp36R2 to generate WT (327 bp) or floxed (514 bp) amplicons (Sup. Fig. 1B).

Recombination of the Pten allele was examined using the primer pair PtenDeltaF1 and PtenDeltaR2 which amplified deleted Pten alleles (397 bp). Recombination of the *Zfp36* allele was examined in multiplex reactions using Zfp36F3, Zfp36R2 and Zfp36R4 primers which amplified the WT (683 bp), floxed (870 bp) or deleted *Zfp36* alleles (769 bp) (Sup. Fig. 1B). All primers are described in Supplementary Table 1.

### Aging GEMM studies

To examine the role of TTP in PCa driven by Pten loss, male experimental mice were generated and allowed to age to 8 weeks, 18 weeks, 38 weeks, or until an ethical endpoint was reached, as determined by Dana-Farber Cancer Institute IACUC and veterinarian stipulations. All aging mice were monitored daily for signs of declining health and manually palpated for prostate tumor development once a week until tumors were detected and then daily following palpation of a prostate tumor. For castration studies, mice received surgical or sham castration at 38 weeks of age and allowed to progress to 50 weeks of age or until a humane endpoint was reached. Following necropsy, whole genitourinary (GU), whole prostate and individual prostate lobe weights were measured, and tissues collected. Pelvic lymph nodes, kidney, liver, and lung tissue samples were collected for assessment of metastatic dissemination. Tissue samples were collected for paraffin-embedded blocks and, where possible, digested and stored in cryopreservation media (10% DMSO in Advanced DMEM/F-12) or snap frozen.

### GEMM tumor RNA-seq analysis

The quality of sequenced reads from the RNA-seq data was assessed, and low-quality reads were filtered using the FastQC tool (Babraham Bioinformatics, Cambridge, UK). Sequence alignment and quantification were performed using the STAR-RSEM pipeline (49, 50). Reads overlapping exons in the annotation of Genome Reference Consortium Mouse Build 38 (GRCm38) were identified. To reduce systemic bias between samples, the Trimmed Mean Method (TMM) was applied to gene level expression counts (51). Genes were filtered out and excluded from downstream analysis if they failed to achieve raw read counts of at least 2 across all the libraries. For the functional enrichment analysis, we used gene set enrichment analysis (GSEA) software (52). Briefly, 1) the Hallmark and Gene Ontology Biological Process gene sets were obtained from MSigDB (53), 2) Log2-median-ratio was used to order genes in the data in a descending manner, 3) Enrichment score (ES) was computed using a Kolmogorov-Smirnov running sum statistics for each gene set, and 4) Significance of the ES was computed using a distribution of null hypothesis which was generated by doing 1,000 random permutations.

### CRISPR-knockout cell lines and RNA-seq

For CRISPR/Cas9-mediated knockout cell line generation, guide RNA (gRNA) sequences CATGACCTGTCATCCGACCA, AAGCGGGCGTTGTCGCTACG and GAGCTCGGTCTTGTATCGAG targeting murine *Zfp36* and human *ZFP36* respectively, were cloned into the lenti-CRISPR/Cas9v2 vector (Addgene, #52961) according to the Zhang lab protocol (54). The scrambled gRNA sequence CACCGCGTGATGGTCTCGATTGAGT was used as a negative control. Viral infection was performed as described by the RNAi consortium (Broad Institute) laboratory protocol “Lentivirus production of shRNA or ORF-pLX clones,” and single clones were isolated following puromycin selection (2 μg/mL; Sigma-Aldrich #P8833). RNA-seq data were aligned with STAR to mouse reference genome mm10 (GRCm38), quantified using RSEM and normalized. Gene set enrichment analysis (GSEA) was performed in GenePattern (genepattern.org). Genotypes were compared as described, using 10,000 gene set permutations to generate normalized enrichments scores, with FDR q-value <0.25 considered significant.

### Immunohistochemical and immunofluorescent staining and quantification of GEMM tissues

For all staining, 4 μm thick sections were cut from paraffin-embedded blocks and dried onto positively charged microscope slides. All antibodies used are listed in Supplementary Table 2.

For immunohistochemistry (IHC), tissue sections were stained using the ImmPRESS® HRP Anti-Mouse IgG (Peroxidase) Polymer Detection Kit (Vector Laboratories) as previously described (55). Tissue sections were imaged using an EVOS FL Auto 2 Cell Imaging microscope (Thermo Fisher Scientific). Images were de-identified and 50 random fields per section were run through QuPath image analysis software (56) to quantify positive DAB-stained cells (as a percentage of total cells).

For immunofluorescence (IF), tissue sections were prepared using standard deparaffinization, rehydration and sodium citrate antigen-retrieval methods. Sections were blocked with 5% goat serum in PBST for 1 hour at room temperature, followed by incubation with primary antibody diluted in 5% goat serum/PBST overnight at 4°C. Where a conjugated primary antibody was used, slides were washed in PBST (4 x 2 minutes) and coverslipped with Vectashield DAPI mounting medium. Where a non-conjugated primary antibody was used, slides were washed in PBST (4 x 2 minutes) and blocked with 5% goat serum/PBST for 15 minutes at room temperature. Sections were incubated with the appropriate fluorescent secondary antibody diluted in 1% goat serum/PBST for 1 hour at room temperature then washed in PBST (4 x 2 minutes). Slides were coverslipped with Vectashield DAPI mounting medium (Vector Laboratories). For analysis, 20 representative images from each tissue section were taken using an EVOS FL Auto 2 Cell Imaging microscope. Staining intensity and area was scored using analysis pipelines generated for CellProfiler software (57).

For Trichrome Staining, tissue sections were stained using the Trichrome Stain Kit (Connective Tissue Stain) (Abcam, ab150686) according to manufacturer’s instructions. Tissue sections were imaged using an EVOS FL Auto 2 Cell Imaging microscope. Images were de-identified and 50 random fields per section were run through QuPath image analysis software (56) to quantify positive reactive stroma-stained area (as a percentage of total cells analyzed).

### Generation of GEMM-derived 2D and 3D cell lines

Organoid (3D) models were generated from GEMM prostate tissue using previously described methods (58). To generate 2D cell lines, organoid cultures were allowed to grow as a single layer in 2D in standard organoid culture medium. Over multiple passages, organoid culture medium was replaced in a stepwise process with standard DMEM supplemented with 10% FBS and passaging until phenotypical and growth rate stability were achieved.

### Organoid budding assay

To prepare for assays, organoid cultures were seeded as single cells in 40 mL Matrigel droplets in 6-well tissue culture plates and cultured for 2 days at 37°C to allow organoid formation. After organoid formation, media was aspirated and Matrigel droplets were manually detached from tissue culture plates and incubated with dispase II (Thermo Fisher, 17105041) at a final concentration of 1-2 mg/mL in Advanced DMEM/F12 (Thermo Fisher, 12634028) for 30 min with gentle shaking at 37°C to remove whole organoids from the Matrigel. Organoids were gently washed twice with PBS and resuspended in 30 mL 33% Matrigel (in Advanced DMEM/F12) at a concentration of 10-20 organoids per well in 96-well tissue culture plates. Starting 24 hrs after replating, daily brightfield images of each organoid were taken using an EVOS FL Auto 2 Cell Imaging microscope. Organoid budding was assessed using an image analysis pipeline generated for CellProfiler software (57) to identify the organoid area deviating beyond the primary sphere shape.

### 2D wound healing assay

GEMM-derived 2D cells (from PB-Cre4/Pten^f/f^ and PB-Cre4/Pten^f/f^/*Zfp36*^f/f^ mice, as well as previously described Pten/Rb1-null murine cell line (48)), were seeded into 24-well tissue culture plates such that they would reach confluence after 48 hrs. Once the cultures were confluent a wound was generated by drawing a sterile p1000 tip gently across the center of each well. Each well was gently washed with PBS to remove all loose cells and debris. Tissue culture plates were subsequently imaged daily using an EVOS FL Auto 2 Cell microscope and wound healing was assessed using the ImageJ Wound Healing Assay macro.

### Organoid growth assessment

For assays, organoid cultures were sparsely seeded sparsely as single cells in 40 mL Matrigel in 96-well tissue culture plates and cultured for 2 days at 37°C to allow organoid formation. After organoid formation, daily brightfield images of individual organoids were taken using an EVOS FL Auto 2 Cell Imaging microscope. Organoid area was measured from images using CellProfiler software (57) and organoid growth assessed as a daily fold-change in organoid area relative to day 0. For each cell line and treatment condition 10 individual organoids were measured.

### Organoid therapy assay

Organoids were established as above for growth assessment assays. Once formed, organoids treated with vehicle (0.1% DMSO), 10 µM enzalutamide (MedchemExpress), 5 µM dimethylaminoparthenlide (DMAPT) or the combination of enzalutamide and DMAPT for 72 hrs. After treatment, cells were incubated with ReadyProbes Cell Viability Imaging Kit Blue/Green (Invitrogen) per well for 30 minutes at room temperature and z-stack images of stained cells were taken using an EVOS FL Auto 2 Cell Imaging microscope. The percentage of cell death was calculated by identifying the percentage of non-viable cells per organoid in at least 10 organoids for each treatment condition.

### *In vivo* therapy experiment

Experiments were carried out on 10-week-old male C57BL/6 mice (The Jackson Laboratory). Allograft models were generated by subcutaneous injection of 2 × 10^6^ GEMM-derived 2D cells (from PB-Cre4/Pten^f/f^ and PB-Cre4/Pten^f/f^/*Zfp36*^f/f^ mice) per animal in 100 µL Matrigel (50% in PBS). Tumors were allowed to establish and grow to approximately 100 mm^3^ before being randomly allocated into vehicle (water), surgical castration, DMAPT (100 mg/kg/day by oral gavage) or castration plus DMAPT combination. All non-castrated treatment groups also received a sham-castration. Tumor volume and animal weight were measured every two days. Tumor volume was measured by caliper and expressed in mm^3^ (tumor volume = 0.5(a x b^2^), where a and b represent the long and short diameter, respectively). Treatment toxicities were assessed by body weight, decreased food consumption, signs of dehydration, hunching, ruffled fur appearance, inactivity, or nonresponsive behavior.

### RT–qPCR

The qPCRs were performed in accordance with MIQE guidelines (59). RNA was harvested from using a standard TRIzol (Thermo Fisher, 15596018) protocol according to the manufacturer’s instructions. Complementary DNA was synthesized using the High-Capacity cDNA Reverse Transcription Kit (Thermo Fisher, 4368813), according to the manufacturer’s instructions. The iTaq™ Universal SYBR® Green Supermix (Bio-Rad, 1725120) was used for PCRs with the cycling conditions recommended in the manufacturer’s instructions. The primers used are detailed in Supplementary Table 1.

### Western Blot

Sub-confluent treated cells were washed twice with cold PBS, trypsinized and then lysed in Pierce RIPA buffer (Thermo Fisher Scientific, 89900) with PhosSTOP™ inhibitor cocktail (Sigma Aldrich, PHOSS-RO) at 4°C for 30 mins. Protein concentrations were measured by a Pierce BCA Protein Assay Kit (Thermo Fisher Scientific, 23225). Proteins were separated by SDS-PAGE (10% Mini-PROTEAN® TGX™ Precast Gel, Biorad, 4561036) and transferred to a PVDF membrane (Bio-Rad, 1620177). The membrane was blocked 3% BSA in TBST for 1 hour at room temperature and then blotted with primary antibodies overnight at 4°C. Full list of antibodies available in Supplementary Table 2. Blots were washed 3 x 5 minutes with TBST. The blots were then incubated with fluorescent-conjugated secondary antibody (Bio-Rad) 1 hour at room temperature and washed 3 x 5 minutes with TBST. Proteins were visualized with a ChemiDoc MP fluorescent imager (Bio-Rad). Bio-Rad Image Lab analysis software was used to quantify protein band density. All western blots were repeated once.

### Statistical analysis

Statistical significance was determined by performing an unpaired two-tailed Student t-test unless otherwise noted. Comparisons between treatment arms in the Kaplan Meier analysis of the *in vivo* study were carried out using a Log-rank (Mantel-Cox) test to compare time-to-ethical-endpoint. A p-value of less than 0.05 was considered as significantly different from the null hypothesis across all experiments as indicated by asterisks in all figures. All statistical calculations were carried out in Prism v9.0 (GraphPad Software, San Diego, CA) and Stata v15.1 (StataCorp, College Station, TX).

## Results

### Loss of *ZFP36*/TTP in prostate cancer patients selects for aggressive disease

In previously published, independent cohorts of localized hormone-sensitive PCa (HSPC), meta-analysis identified a consistent association with *ZFP36*/TTP loss and increased risk of biochemical recurrence and development of lethal disease after curative therapy (Fig. 1A, left) (14, 37). *ZFP36*/TTP loss was defined by expression below the cohort median. We then developed a *ZFP36* loss signature (Fig. 1B) which was applied to whole transcriptome datasets with associated clinical outcomes and a consistent prognostic effect was again observed (Fig. 1A, left lower). To confirm that TTP protein expression holds a similar association, a tissue microarray was created from a cohort of men with localized PCa undergoing radical prostatectomy (RP) and fluorescent IHC for TTP was performed on RP specimens (Fig. 1C). Loss of TTP protein expression was significantly associated with shorter disease-free survival, a finding consistent with a previously reported independent cohort (Mahajan *et. al.* (37). The combined results are reported as pooled meta-analysis (Fig. 1A, right upper). We further documented in a meta-analysis of published data (14) that localized PCa with *ZFP36* RNA levels in the lower quartile was associated with an almost 2-fold risk of lethal PCa (metastatic disease or death from PCa) (Fig. 1A, bottom right).

**Figure 1:**
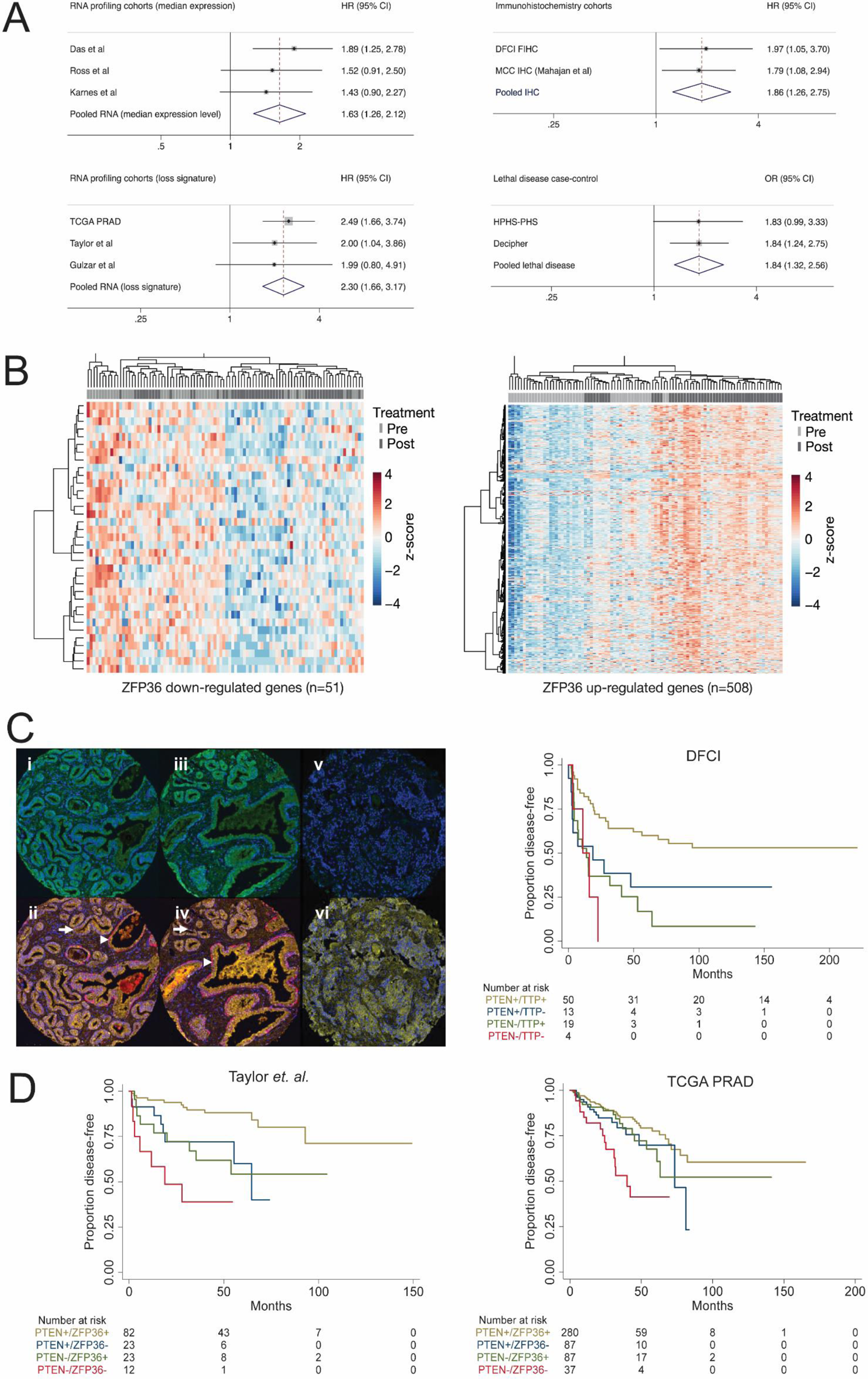
*ZFP36*/TTP and clinical outcomes. (A) Forest plots depicting RNA and IHC-based for *ZFP36*/TTP expression related to clinical outcomes (biochemical recurrence and disease-free survival) and risk of lethal PCa (case-control cohorts). (B) (left) Upregulated and downregulated genes were identified by differential expression analysis of TCGA PRAD cases divided by lower quartile expression of *ZFP36*; (right). (C) Representative images of IF staining in human PCa used for expression analysis. Benign glands (arrowheads) stain for pan-cytokeratin (yellow) and basal (red) cocktails; tumor cells (arrows) demonstrate absent basal expression (panels ii & iv). Corresponding sections (i & iii) demonstrate intact epithelial staining for TTP (green). Panels v-vi: Diffuse prostate tumor with absent TTP expression. (D) Kaplan Meier survival analysis demonstrating that TTP deficiency, measured by protein expression (DFCI (39, 40)) and *ZFP36* mRNA expression (TCGA PRAD (31); Taylor et al (36)), results in shorter disease-free-survival, and even shorter disease-free-survival in combination with PTEN deficiency.

Given the literature that cooperative loss of two or more tumor suppressor genes can drive more aggressive disease (48, 60, 61), we explored the proposition that low expression of TTP increases the aggressiveness of tumors with PTEN loss. We performed additional IHC for PTEN and observed that men within the lower quartile expression of loss of one tumor suppressor had a shorter disease-free survival (PTEN low - HR: 2.86, 95% CI 1.61-5.10, p = 0.0002; TTP low - HR: 1.97; 95% CI 1.05-3.70, p = 0.03) (Fig. 1D left panel). Notably, men with both low TTP and low PTEN expression had the poorest disease-free survival (HR 2.98, 95% CI 1.05-8.46, p = 0.03), which remained significant in multivariable analysis adjusting for prognostic clinicopathological factors of Gleason score, T stage and PSA (HR: 4.50, 95% CI 1.03-19.61, p = 0.045). The finding that lower levels of TTP and PTEN portend the highest risk of relapse was independently supported by gene expression analyses from two independent cohorts (31, 36) noting patients with transcript levels of *ZFP36* and *PTEN* alone in the lower quartile had poorer disease-free survival than men with higher levels of both *ZFP36* and *PTEN,* and that men with concurrent low levels of *ZFP36* and *PTEN* had the highest risk of relapse (Fig. 1D). These data validate in multiple independent datasets that low TTP/*ZFP36* expression is related to poor prognosis in PCa, and that the additional loss of PTEN expression corresponds with even more aggressive disease.

### Tristetraprolin and *ZFP36* expression is upregulated in prostate tumors in response to enzalutamide treatment

To examine the clinical response of *ZFP36*/TTP to enzalutamide treatment we examined two clinical cohorts where the prostate tumors from the same patient were investigated pre- and post-enzalutamide treatment. PCa samples from the DARANA study (Dynamics of Androgen Receptor Genomics and Transcriptomics After Neoadjuvant Androgen Ablation (41)) displayed significant upregulation of *ZFP36*/TTP at the protein (p < 0.0001, Fig. 2A) and gene expression (p = 0.0001, Fig. 2B) level following enzalutamide treatment. Similar RNA *ZFP36* upregulation was observed in an independent neoadjuvant ADT study - the NCI dataset of paired pre- and post-treatment PCa samples treated with neoadjuvant ADT plus enzalutamide (p < 0.0001, Fig. 2B). In the NCI tumor dataset, we examined the correlation between *ZFP36* expression and the volume of post-treatment residual tumor (Fig. 2C). *ZFP36* expression pre-treatment did not correlate with the volume of post-treatment residual tumor (Spearman R-value = -0.2187, p = 0.2). However, there was a significant negative correlation in *ZFP36* expression post-treatment samples (Spearman R-value = -0.3566, p = 0.033), indicating that the largest residual tumors post-enzalutamide occurred in patients where *ZFP36* RNA expression was low pre- and post-treatment. Following enzalutamide treatment, patient samples from the DARANA study showed significant increases in H3K27acetylation at the *ZFP36* locus post-enzalutamide (Fig. 2D-E, p = 0.0003), epigenetically confirming enhanced transcriptional activation of this locus following treatment.

**Figure 2:**
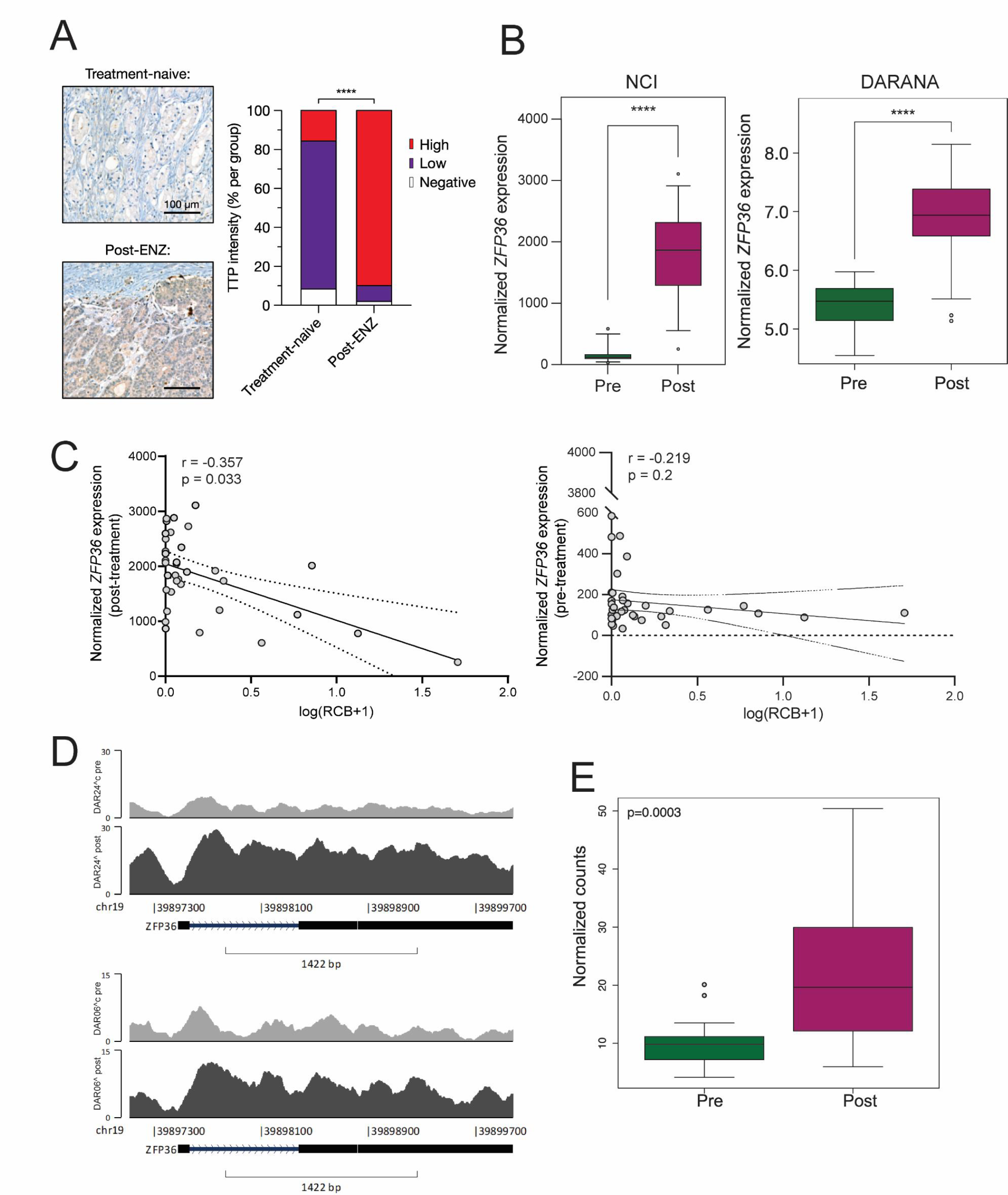
*ZFP36*/TTP expression in response to enzalutamide. (A) Representative images and quantification of TTP IHC staining intensity in DARANA patient tissues comparing treatment-naïve and post-enzalutamide samples. (B) *ZFP36* expression from RNA-seq pre- and post-enzalutamide in the NCI and DARANA clinical studies. (C) Correlation of normalized *ZFP36* expression vs volume of post-treatment residual cancer burden (RCB) in pre- and post-enzalutamide samples (n = 36 patients). (C) Representative H3K27 acetylation tracks at the *ZFP36* locus from two DARANA patients, comparing pre- and post-enzalutamide samples, and (E) quantification of H3K27 acetylation signal at the *ZFP36* locus pre- and post-enzalutamide treatment.

### Loss of *Zfp36* accelerates disease progression in a mouse model of prostate adenocarcinoma driven by Pten deletion

To model the clinical finding that low expression of *ZFP36* synergizes with *PTEN* loss to drive aggressive PCa progression, we engineered *Zfp36* deletion in a previously characterized mouse model of prostate adenocarcinoma induced by *Pten* deletion (62). Our model utilized expression of the PBCre4 transgene (47) to ensure prostate epithelial exclusive deletion of *Zfp36* and *Pten* floxed alleles (Supp. Fig. 1A). PBCre4:*Pten*^f/f^ (*Pten^f/f^)* mice develop PIN by 6-8 weeks and progress to invasive adenocarcinoma with rare incidence of metastatic disease (62) (Fig. 1B). In parallel, PBCre4:*Zfp36*^+/f^ (*Zfp36*^+/f^) and PBCre4:*Zfp36*^f/f^ (*Zfp36*^f/f^) mice developed PIN but did not progress to adenocarcinoma (Supp. Fig. 2A). The effect of *ZFP36* loss alone was also assessed *in vitro* utilizing a primary human prostate epithelial model immortalized by expression of hTERT with co-expression of AR (957E/hTERT, PrEC-AR) (63). Loss of *ZFP36* expression in this model resulted in overall increased cell proliferation (Supp. Fig. 2B). Notably, this model showed that *ZFP36* loss increased proliferation. With *ZFP36* intact, the cells exhibit the previously reported androgen (R1881)-mediated anti-growth phenotype (63), which was significantly attenuated by *ZFP36* loss (Supp. Fig. 1B). Further examination of these genotypes using GEMM-derived 2D cell lines demonstrated only homozygote loss of *Zfp36* enabled cell transformation and growth, equivalent to that of organoids exhibiting *Pten* loss (Supp. Fig. 2C). Together, these data clearly demonstrate that *ZFP36* can reprogram AR signalling and induce PIN, a precursor lesion to primary prostate adenocarcinoma.

Histological analyses of collected prostates from mice either harbouring prostate specific loss of *Pten* alone or in combination with heterozygous or homozygous loss of *Zfp36* clearly indicated the loss of *Zfp36* significantly increased progression of PCa initiation and progression of primary disease (Fig. 3A). The combined weights of the dorsolateral and ventral prostate lobes in mice with combined *Pten* and *Zfp36* loss (PBCre4:*Pten^f/f^*:*Zfp36^+/f^*, 190.0 ± 53.5 mg; PBCre4:*Pten^f/f^*:*Zfp36^f/f^*, 229.8 ± 71.1 mg) was significantly increased compared to *Pten^f/f^* mice (118.8 ± 43.7 mg) at 18 weeks. This increase in combined dorsolateral and ventral prostate lobe weight was also present at 38 weeks of age (Fig. 3B). Accordingly, mice with combined *Pten* and *Zfp36* loss exhibited significant shorter median survival of 53.4 weeks for *Pten^f/f^*:*Zfp36^+/f^* and 49.6 weeks for *Pten^f/f^*:*Zfp36^f/f^* mice compared with 66.9 weeks for *Pten^f/f^* mice (p < 0.0001) (Fig. 3C).

**Figure 3:**
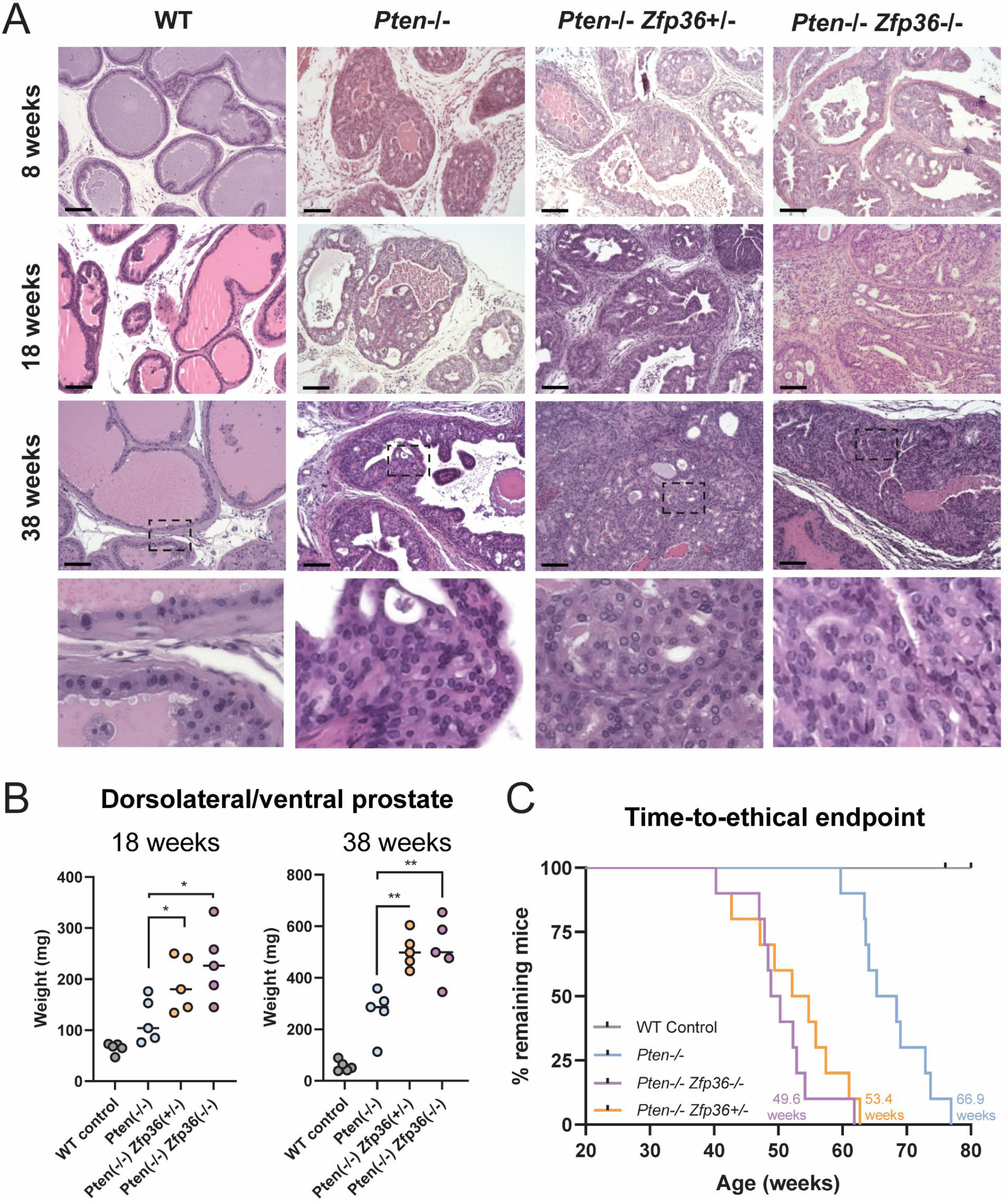
*Zfp36* loss accelerates progression of prostate cancer in Pten-null murine tumors. (A) hematoxylin and eosin staining of murine tumors highlighting morphological progression of wild-type (WT), Pten^f/f^/*Zfp36*^+/+^ (*Pten*-/-), Pten^f/f^/*Zfp36*^f/+^ (*Pten*-/- *Zfp36*+/-) and Pten^f/f^/*Zfp36*^f/+^ (*Pten*-/- *Zfp36*-/-) dorsolateral prostate tissue at 8, 18, and 38 weeks. Scale bar 100 µm. (B) Comparative weight of dorsolateral and ventral prostate tissue in GEMMs at 18 and 38 weeks. (C) Kaplan Meier graphs from GEMM aging studies show that prostate-specific deletion of *Zfp36* significantly reduces time-to-ethical endpoint in PCa driven by loss of Pten. *p<0.05, **p<0.005.

To further characterize changes mediated by *Zfp36* loss, RNA-seq from mouse tumors was performed. Given the molecular regulation of NF-κB activity by *Zfp36* (14) it was not surprising that GSEA Hallmark analysis indicated significant enrichment for inflammatory associated gene signatures (Fig. 4A, Supp. Table 3). GSEA Hallmark analysis data comparing *Pten^f/f^*:*Zfp36^f/f^ vs. Pten^f/f^* GEMM prostate tumors was strikingly similar to that of DARANA clinical data comparing pre-vs. post-enzalutamide prostate tumors (Fig. 4A). Further validation of NF-κB activation (p-p65) and overall stromal inflammation was performed by IHC on tumors extracted from GEMMs at 38 weeks (Fig. 4B). These data confirmed that the observed aggressive nature of prostate tumors harboring *Pten:Zfp36* loss was in part due to a significant increase in tumor cell NF-κB activity and a resulting overall inflammatory phenotype.

**Figure 4:**
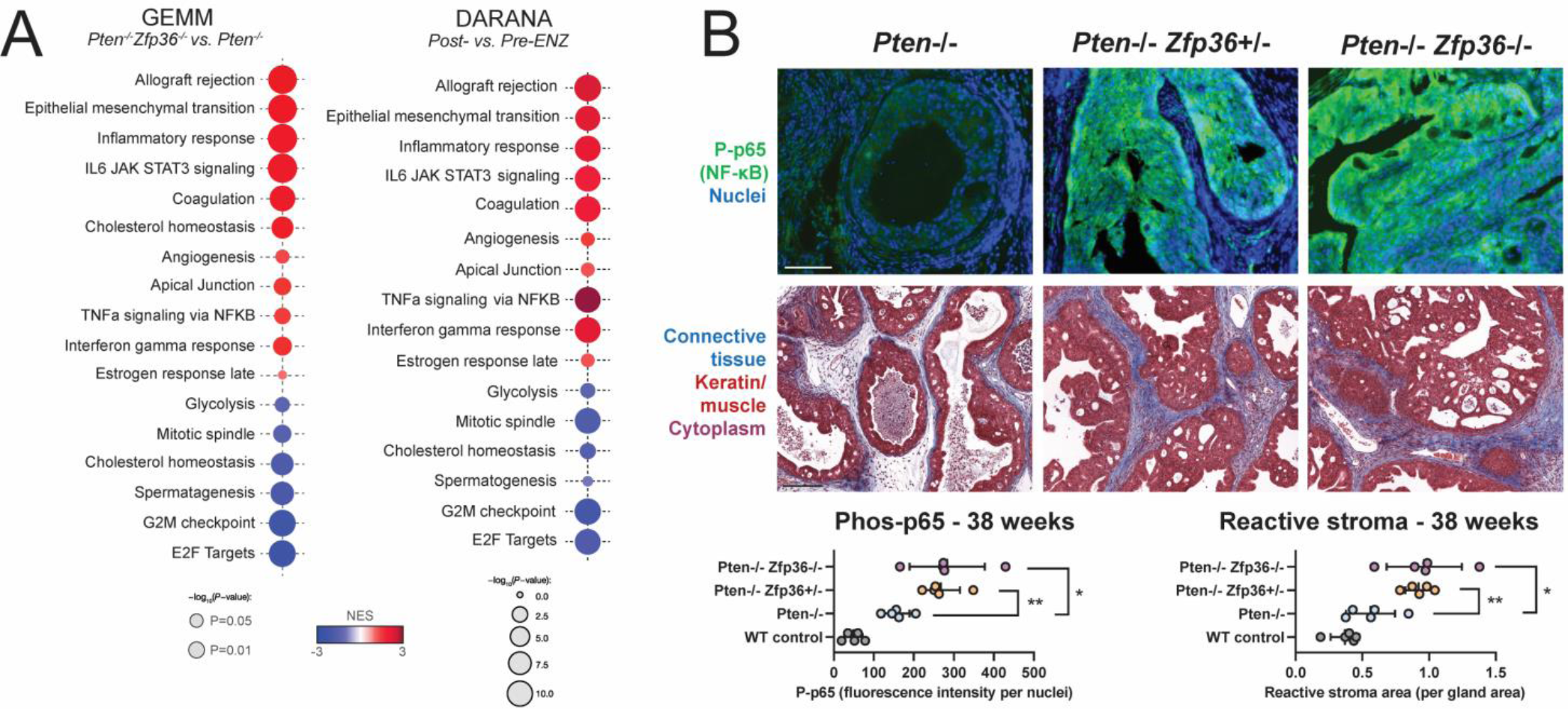
*Zfp36* loss increases an inflammatory prostate cancer phenotype in Pten-null murine tumors. (A) GSEA from RNA-seq of endpoint GEMM PCa tumors comparing *Pten*-/-, and *Pten*-/- *Zfp36*-/- GEMMs, highlighting positively and negatively enriched Hallmark pathways. (B) Phos-p65 IF and Masson’s Trichrome staining PCa in *Pten*-/-, *Pten*-/- *Zfp36*+/- and *Pten*-/- *Zfp36*-/- GEMM dorsolateral prostate tissue at 38 weeks, with corresponding quantification. Scale bar 100 µm. *p<0.05, **p<0.005.

Additional analysis using Gene Ontology Biological Process (GOBP) from our RNA-seq from GEMM tumors identified significant positive enrichment of inflammation and leukocyte gene expression signatures in *Pten^f/f^*:*Zfp36^f/f^* mice compared with *Pten^f/f^* mice (Fig. 5A, Supp. Table 4). Due to this enrichment of immune related gene signatures and concern for an immune infiltrate being the cause for these results we performed additional RNA-seq on *Pten* deleted 2D PCa cell lines generated from our GEMMs (48). We further engineered these cells for *Zfp36* loss by use of CRISPR/Cas9. RNA-seq from these 2D cell lines showed complementary results, indicating that our observed induction of immune related gene signatures was autonomous to the tumor cell (Supp. Table 5-6, Supp. Fig. 3). Leukocyte gene signatures that were enriched in our gene expression data from prostate luminal epithelial cells were related to chemotaxis, proliferation, and migration. We validated these attributes in tumor tissue from mice indicating that loss of *Zfp36* significantly increased overall cell proliferation, in line with increased proliferation in prostate cell lines (Supp. Fig. 2) and invasion potential by degradation of the basement membrane (Fig. 5B). Aligned with this aggressive phenotype was an increased incidence of micro-metastasis to the kidney, liver, lung and macro-metastasis to the pelvic lymph nodes in a subset of mice examined with *Zfp36* loss (Fig. 5C-D). Further validation of this *in vivo* metastatic phenotype was by an indicated increase of invasive potential (organoid budding in soft matrigel) and cell motility in GEMM derived *ex vivo* models (Fig. 5E-F).

**Figure 5:**
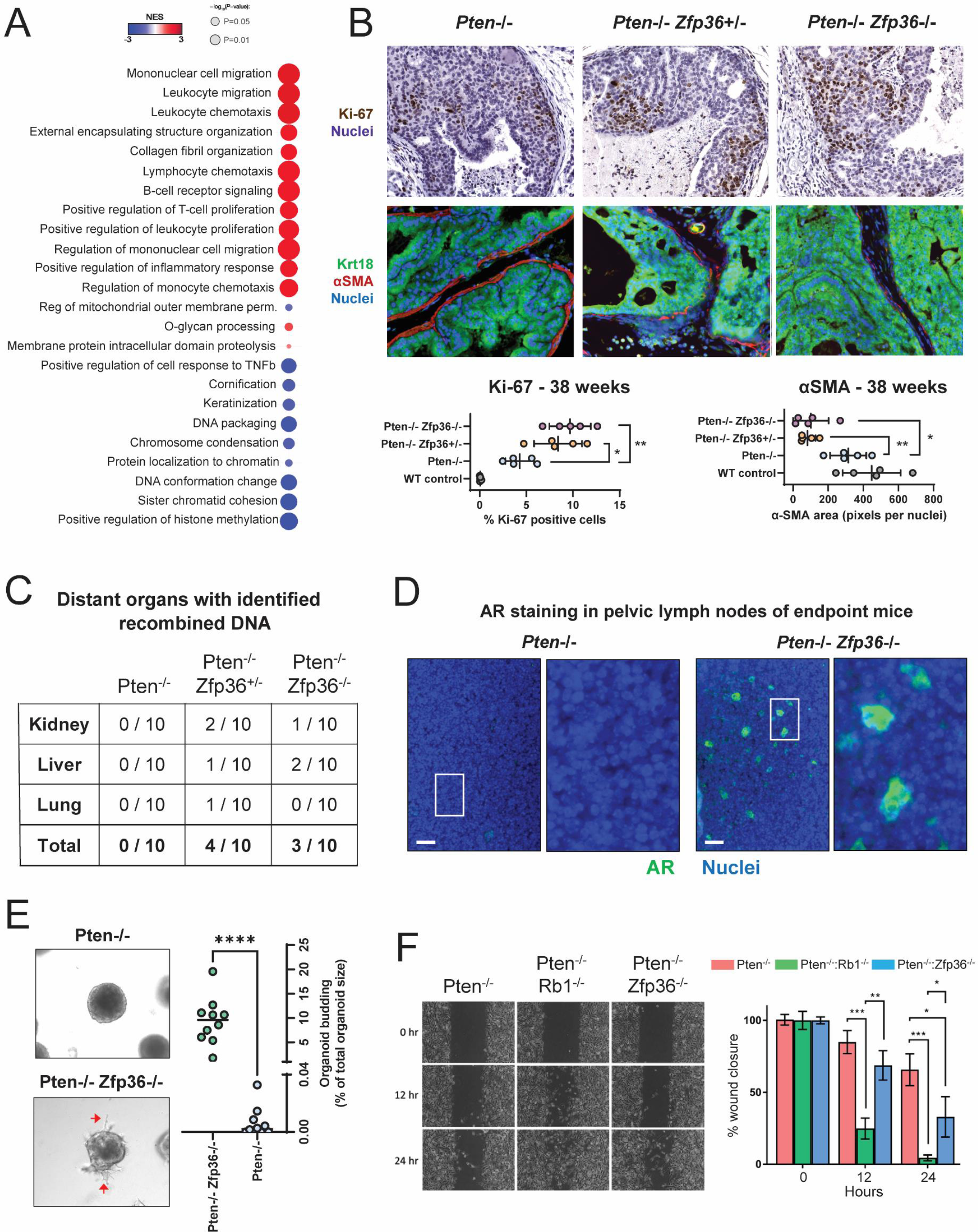
Increased metastatic potential occurs with *Zfp36* loss Pten-null murine tumors. (A) GSEA from RNA-seq of endpoint GEMM PCa tumors comparing *Pten*-/-, and *Pten*-/- *Zfp36*-/- GEMMs, highlighting significant positively and negatively enriched GOBP pathways. (B) Ki-67 IHC, and Krt18 and αSMA IF staining PCa in *Pten*-/-, *Pten*-/- *Zfp36*+/- and *Pten*-/- *Zfp36*-/- GEMM dorsolateral prostate tissue at 38 weeks, with corresponding quantification. Increased tumor cell proliferation and basement membrane breakdown is observed with loss of *Zfp36*. (C) Number of mice that displayed PCa cells in distant organs by recombination PCR in *Pten*-/-, *Pten*-/- *Zfp36*+/- and *Pten*-/- *Zfp36*-/- GEMMs. (D) Representative androgen receptor (AR) IF staining in pelvic lymph nodes of *Pten*-/- and *Pten*-/- *Zfp36*-/- GEMMs highlighting local dissemination of prostate cells. Scale bar 100 µm. AR staining was over exposed during imaging to assist with prostate cell identification. (E) Representative images and quantification of budding in GEMM-derived organoids highlighting increased invasive and metastatic potential of *Pten*-/- *Zfp36*-/- organoids. (F) Scratch assay in GEMM-derived 2D cells, comparing *Pten*-/- and *Pten*-/- *Zfp36*-/- wound healing with that of *Pten*-/- *Rb1*-/-, a previously described metastatic, neuroendocrine PCa murine cell line (48). *p<0.05, **p<0.005, ***p<0.001, ****p<0.0001.

Wound closure of *Pten^f/f^*:*Zfp36^f/f^* cells occurred significantly faster than for *Pten^f/f^* cells, but was slower compared to a previously described metastatic, neuroendocrine PCa model, *Pten^f/f^*:*Rb1^f/f^* (48), which hints that loss of *Zfp36* may result in an intermediate phenotype, poised to become aggressively metastatic.

### *Zfp36* loss induces tumor cell changes associated with phenotypic plasticity

Given the observed rapid disease progression and gene set enrichments following loss of *Zfp36*, we hypothesized that phenotypic plasticity was occurring given similar findings when loss of other tumor suppressors occur with PTEN loss (48, 60). To that end, tumors extracted from *Pten^f/f^*, *Pten^f/f^*:*Zfp36^+/f^* and *Pten^f/f^*:*Zfp36^f/f^* GEMMs at 38 weeks were interrogated by IF staining to examine androgen receptor (AR), synaptophysin (Syp) and CD45 antigen (CD45) expression. Low AR expression with corresponding high Syp expression, is characteristic of neuroendocrine PCa. Tumors from *Pten^f/f^* mice retained AR and had minimal synaptophysin and CD45 expression, while GEMM tumors with combined loss of *Pten* and *Zfp36* displayed an inversed staining pattern that included reduced AR and increased synaptophysin and CD45 in a heterogeneous manner (Fig. 6A). To validate that CD45 staining was predominantly exclusive to luminal prostate cells, as suggested by our gene expression data, we performed co-staining of cytokeratin-8 (Krt8 – a marker for luminal prostate epithelium) and CD45. Fig. 6B provides clear validation that prostate luminal cells from mice with *Pten^f/f^*:*Zfp36^f/f^* deletion do co-express both the luminal marker Krt8 with CD45. These data indicate that *Zfp36/ZFP36* is a critical mediator of prostate luminal cell identity.

**Figure 6:**
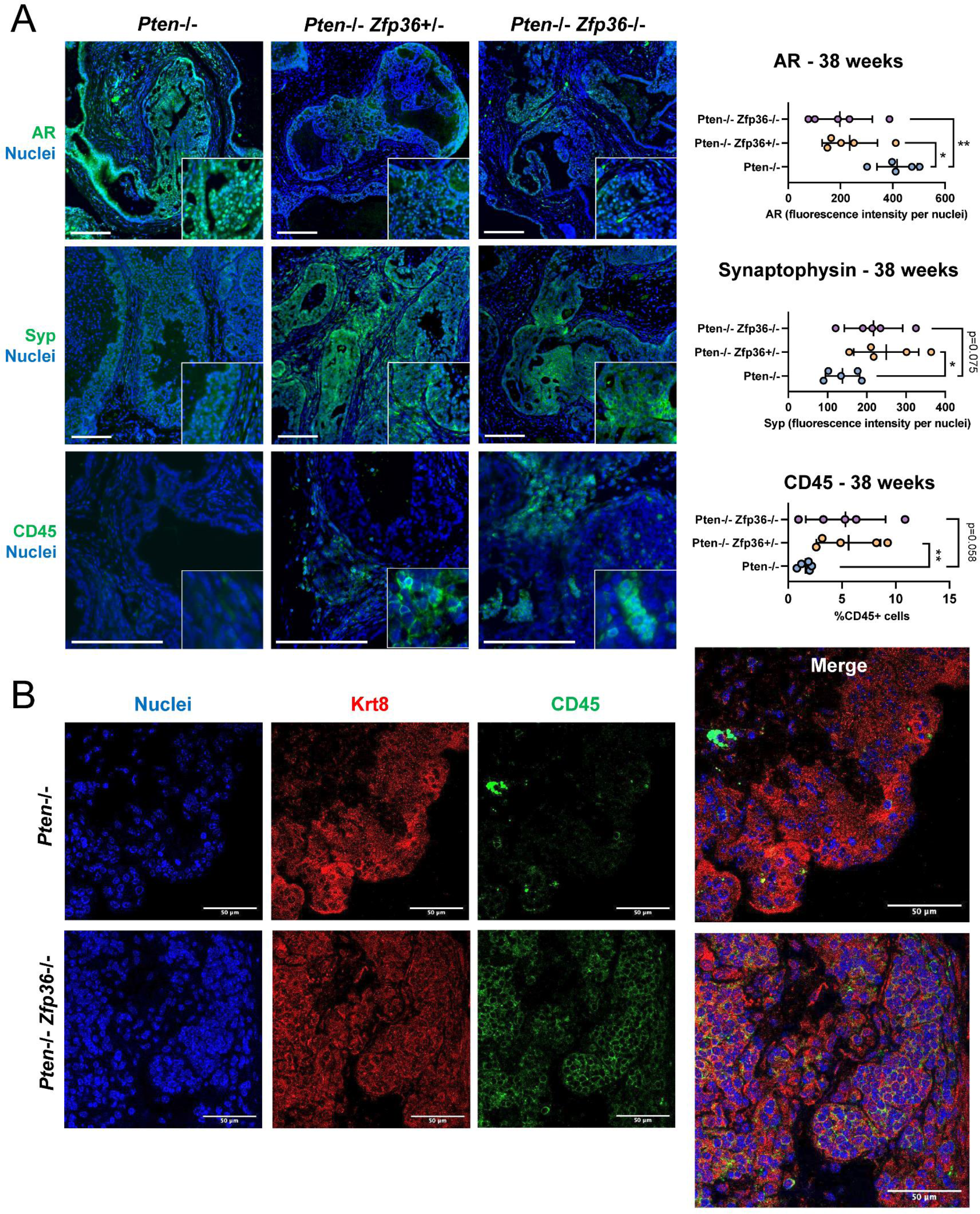
Loss of *Zfp36* induces phenotypic plasticity in Pten-null murine prostate tumors. (A) AR, synaptophysin (Syp) and CD45 IF staining PCa in *Pten*-/-, *Pten*-/- *Zfp36*+/- and *Pten*-/- *Zfp36*-/- GEMM dorsolateral prostate tissue at 38 weeks, with corresponding quantification. Scale bar 100 µm. *p<0.05, **p<0.005. (B) Dual Krt8 and CD45 IF staining in *Pten*-/- and *Pten*-/- *Zfp36*-/- GEMM dorsolateral prostate tissue at 38 weeks. Scale bar 50 µm.

### Loss of *Zfp36* increases resistance to hormonal therapy in Pten-null tumors

Our current results showing *Zfp36* loss lowers AR expression and induces phenotypic plasticity associated with NF-κB activation is demonstrated to drive resistance to androgen deprivation therapy and progression to castration-resistant PCa (11, 48). To test the response to androgen deprivation, we surgically castrated 38-week-old, tumor-bearing *Pten^f/f^, Pten^f/f^*:*Zfp36^+/f^*, and *Pten^f/f^*:*Zfp36^f/f^* mice and monitored their survival. Castration, as expected appeared curative in *Pten^f/f^* mice, whereas castration of *Pten^f/f^*:*Zfp36^+/f^* and *Pten^f/f^*:*Zfp36^f/f^* mice resulted in a median survival of 56.1 and 53.9 weeks respectively (Fig. 7A), indicating little to no effect of androgen deprivation. Notably, there was no observed increased survival benefit following castration in *Pten^f/f^*:*Zfp36^+/f^* and *Pten^f/f^*:*Zfp36^f/f^* mice. Further, a cohort of mice had tumors excised for analysis 12 weeks post-castration and demonstrated a significant increase in prostate weight (Fig. 7A). These initial observations were validated by using GEMM derived *ex vivo* models to transplant to naïve wildtype mice or use as 3D organoid models (Fig. 7B, Supp. Fig. 4A). All *in vitro*, *in vivo* and *ex vivo* models independently demonstrate that loss of *Zfp36* drives rapid acquisition to a castration-resistant phenotype. It had previously been reported that castration-resistance could be driven by phenotypic plasticity by loss of tumor suppressor genes, *Pten, Rb1* and/or *Tp53* (39, 48, 60). Using our GEMM derived *ex vivo* models, we assessed if our observed phenotypic plasticity and resistance to castration driven by *Zfp36* loss was associated with *Rb1* and *Tp53* loss of function. Using the Mdm2 antagonist (or ‘p53 activator’) Nutlin-3 (5 µM) and CDK4/6 inhibitor Palbociclib (2 µM) we show that percent of surviving cells after Nutlin or Palbociclib following *Zfp36* loss does change and thus our described aggressive phenotype mediated through *Zfp36* loss is independent of the functional loss of *Rb1* or *Tp53* (Supp. Fig. 4B).

**Figure 7:**
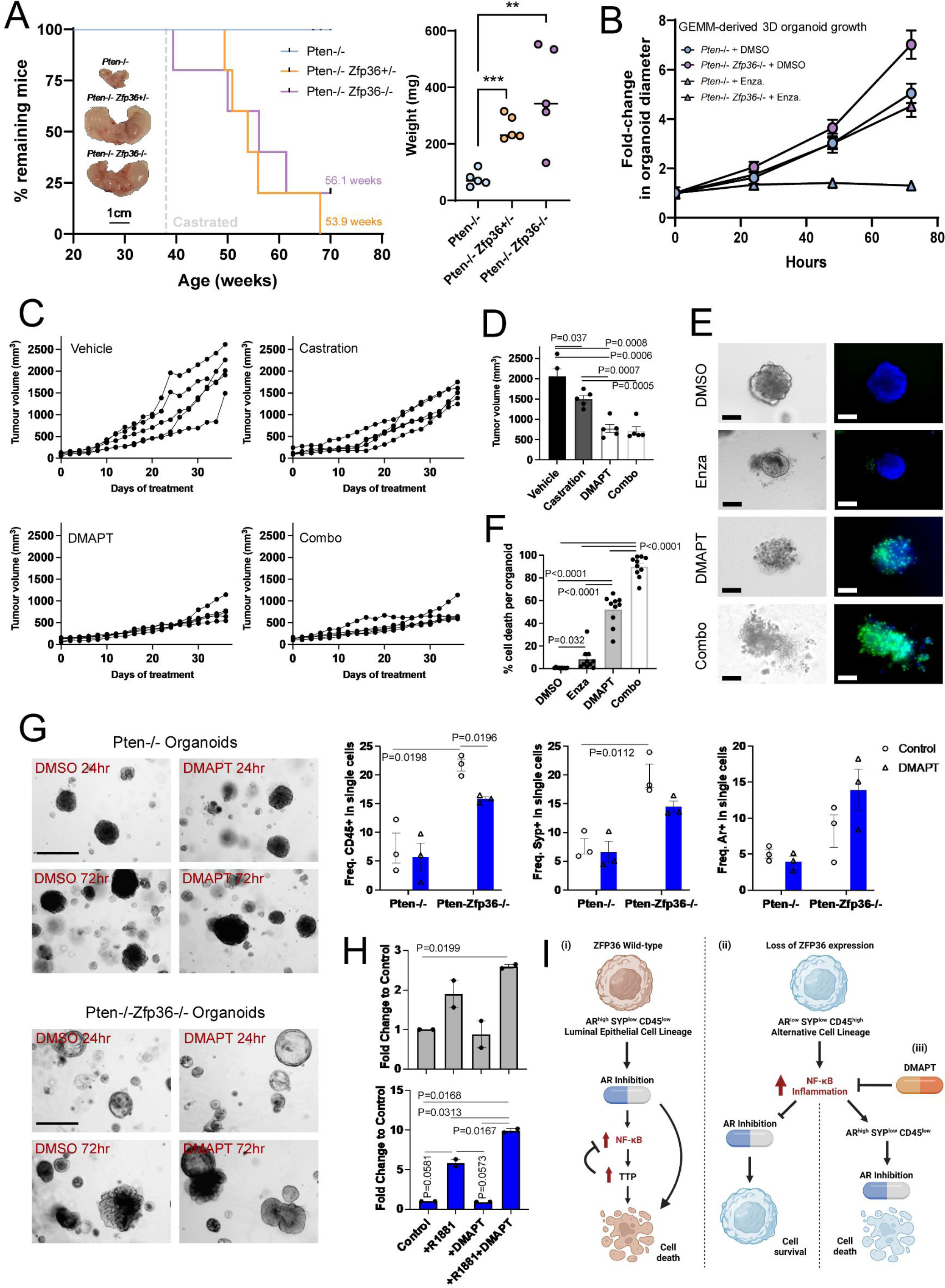
*Zfp36* loss drives castration resistance in Pten-null murine tumors, which is counteracted with the NF-κB inhibitor DMAPT. (A) Kaplan Meier graph from GEMM aging studies where mice were surgically castrated at 38 weeks comparing *Pten*-/-, *Pten*- /- *Zfp36*+/- and *Pten*-/- *Zfp36*-/- mice, and whole prostate weights and representative images from mice 12 weeks post-castration. (B) Quantification of GEMM-derived *Pten*-/- and *Pten*-/- *Zfp36*-/- organoid growth in the presence and absence of enzalutamide (10 µM). (C) Allograft tumor growth in mice treated with DMAPT (100 mg/kg/day) or water vehicle ± surgical castration, n=5 mice per treatment group. (D) End-point tumor volumes from allograft therapy studies. (E) Representative images and (F) quantification of cell death (green) in GEMM-derived PCa organoids treated with DMAPT (5 µM), enzalutamide (10 µM) or the combination of both for 72 hours, n=10 organoids per treatment group. (G) Representative images and flow cytometry quantification for CD45, synaptophysin, and AR expression in GEMM-derived PCa organoids treated with DMAPT (5 µM) or DMSO vehicle for 72 hours. (H) Fold-change in expression of the AR response gene – *Fkbp5* in GEMM-derived 2D cell lines treated with DMAPT (5 µM) or DMSO vehicle for 72 hours, R1881 (10nM was used to stimulate AR activity. *p<0.05, **p<0.005, ***p<0.001. (I) Schematic overview: i) when *ZFP36* is intact epithelial cells present with a luminal lineage phenotype and sensitivity to AR inhibition. In response to increased inflammation and NF-κB expression, ZFP36/TTP increases as part of a negative feedback loop. ii) loss of *ZFP36* results in an alternative epithelial cell lineage phenotype with reduced AR expression, increased SYP and CD45 expression, and uncontrolled NF-κB activation and inflammation, leading to lack of response to AR inhibition. iii) DMAPT treatment inhibits NF-κB and inflammation signaling and restores a more luminal epithelial cell type and responsiveness to AR inhibition.

### TTP loss increases sensitivity of Pten-null tumors to NF-κB inhibition

With the evidence that the phenotypic effects of *Zfp36* loss in Pten-null tumors is associated with increased NF-κB activity we sought to determine the efficacy of an oral NF-κB inhibitor, dimethylaminoparthenolide – DMAPT (11, 64, 65) could 1) demonstrate effect as monotherapy *in vivo* in tumors with *Zfp36* loss, and 2) re-sensitize tumors with *Zfp36* loss to castration. Allograft *in vivo* models were generated by subcutaneously injecting GEMM-derived *Pten^f/f^*:*Zfp36^f/f^* cells into C57BL/6N mice. Surgical castration (1500mm^3^ mean end-tumor volume) had a minimal effect on tumor growth compared to vehicle-treated (2056 mm^3^) mice (Fig. 7C and E). Treatment with DMAPT (100 mg/kg/day) alone (772.5 mm^3^) or in combination with surgical castration (717.9 mm^3^) reduced tumor growth by 63.4-65.1% compared to vehicle. Similarly, *in vitro*, GEMM-derived *Pten^f/f^*:*Zfp36^f/f^* organoids showed a significant cell death response to DMAPT (5 µM) both alone and in combination with enzalutamide (10 µM) (Fig. D and F). To confirm the specificity of DMAPT antitumor effect is dependent on NF-κB activation, silencing p65-NF-κB in *Pten^f/f^*:*Zfp36^f/f^* cells significantly reduced response to DMAPT with concurrent sensitization to enzalutamide (Supp. Fig. 5). In addition to improving control of tumor growth and response to AR inhibition, treating *Pten^f/f^*:*Zfp36^f/f^* PCa organoids with DMAPT partially reversed the phenotypic effects of *Zfp36* loss. This included reduction of CD45 and synaptophysin expression and increased AR expression and function (Fig. 7G-H). Jointly, these data confirm pharmaceutical inhibition of NF-κB as a viable strategy to treat and revert therapy resistance in PCa.

## Discussion

Metastatic PCa remains lethal despite significant extension of overall survival with the addition of docetaxel, abiraterone, apalutamide or enzalutamide to testosterone suppression (66–70). We have previously demonstrated that combinatorial loss of tumor suppressor genes (TSG’s) PTEN, RB1 and/or P53 drive phenotypic plasticity, metastatic progression, and resistance to ARSI (48, 61). Moreover, single-cell RNA-seq analysis using these mouse models of phenotypic plasticity (48) suggests that the induction of inflammatory response pathways is associated with multilineage potential of tumor cells (71). In the context of hormone-sensitive metastatic PCa PTEN is a well described tumor suppressor gene that occurs early in about 40% of patients (39, 72). Based on the premise that concurrent loss of 2 or more TSG’s including PTEN, P53, and RB1 is associated with more aggressive metastatic PCa (48, 60), and P53 and RB1 loss are often late events first noted in castration resistant disease, we asked if an alternate route of phenotypic plasticity could be activated in the context of PTEN loss and potentially independent of P53 and RB1 loss.

Resistance to androgen receptor signaling inhibition (ARSI) can be mediated by both nuclear NF-κB localization/activation (73, 74) and phenotypic plasticity with neuroendocrine differentiation (75, 76). Both events predict poor prognosis for patient’s but their precise contribution to PCa progression is unknown. We recently demonstrated that neo-adjuvant ARSI neo-adjuvant treatment of high-risk PCa patients resulted in significant enrichment of NF-κB signaling and a gene signature associated with neuroendocrine-like disease state (41). Also, we previously identified *ZFP36* as a key regulator of NF-κB signaling (14) and as such hypothesized that *ZFP36* loss may drive PCa progression by creating a pro-proliferative, inflammatory tumor environment, and features of phenotypic plasticity. In recent years, independent research groups have suggested that *ZFP36* functions as a tumor suppressor since its expression is suppressed in various tumor types compared with normal tissues. In both breast and PCa, low *ZFP36* expression is a negative prognostic indicator, and it is associated with rapid tumor progression and poor clinical outcome (16, 77). Here, we confirmed that *ZFP36* is biologically relevant in PCa, resulting in increased tumor progression and resistance to ARSI in PTEN-null patients and tumor models. Additionally, loss of *ZFP36* in normal epithelial cells results in cell transformation significant enough to develop prostatic hyperplasia and PIN, a precursor to prostate adenocarcinoma. This implies that a ‘second hit’, such as loss of *PTEN*, appears to be required to drive prostate cells to cancer. We have previously reported upregulation of NF-κB/p65 in PCa following castration (11) and analysis of DARANA patient tumors (Fig. 4A) confirmed increased activation of inflammatory response genes, including NF-κB in post-enzalutamide samples. In fact, TNFα signaling via NF-κB was the top positive enrichment from RNA-seq of post-enzalutamide tumor samples (Fig. 4A). The matching upregulation of *ZFP36*/TTP in these post-enzalutamide tumors (Fig. 2) clearly highlights the functional role of TTP to negatively regulate NF-κB-driven inflammation (Fig. 7I). Examination of the NCI neoadjuvant ADT tumor dataset demonstrates that *ZFP36* expression is not only elevated upon enzalutamide treatment, but also differs between responders and non-responders (Fig. 2C). Despite a global increase in *ZFP36* expression, the lowest levels of *ZFP36* activation post-treatment were observed in patients with the greatest residual tumor burden, highlighting the significant effect that functional TTP appears to be having on response to therapy. In the absence of functional TTP (in GEMM and GEMM-derived models) inflammation is uncontrolled and GSEA Hallmark analysis looks remarkably like that of post-enzalutamide tumors (Fig. 4A).

Where tumor development was already being driven by loss of the tumor suppressor PTEN, additional loss of *ZFP36* resulted in pronounced increase in PCa progression *in vivo*. While *Pten^f/f^*:*Zfp36^+/f^* and *Pten^f/f^*:*Zfp36^f/f^* mice did not develop lethal macro-metastatic disease, but rather local and distant dissemination (micro-metastasis) was observed in a subset of mice harboring *Zfp36* loss. We attribute this lethal phenotype, at least partially, to the induction of phenotypic plasticity. While the enrichment of nervous system development gene sets is noted, inflammation and leukocyte cell identity and function gene sets appear more significantly enriched, highlighting the ability of PCa cells to activate a process termed lymphocyte mimicry. Cancer cell lymphocyte (immune) mimicry has recently been described as a distinct cancer hallmark (78) and describes tumor cells taking on immune-like genetic profiles or bulk tumors trafficking in greater numbers of immune cells (79, 80). Our data demonstrates activation of lymphocyte mimicry by loss of *Zfp36* is tumor cell autonomous, supporting the concept of tumor cell phenotypic plasticity. It had been previously noted that lung cancer cells could activate a lymphocyte mimicry program by adoption of gene activation normally restricted to lymphocytes. These attributes were noted to include anchorage independent mobilization, chemokine response, and modulation of local inflammatory conditions. It was observed that these lung cancer cells gained this ability via the upregulation of the lymphocyte restricted transcription factor and chromatin regulator, Aiolos (gene *IKZF3*) (81). It was later shown that Aiolos cooperated with STAT3 to drive chemokine receptor expression and promote breast cancer metastasis (82). Further, lymphocyte mimicry has also been attributed in cancer cells by chromosome instability (CIN). Downstream of CIN, it was shown that chronic activation of cytosolic DNA sensing pathways including STING, resulted in upregulation of interferon signaling and NF-κB activity. Instead of a lethal response by epithelial cells, it was instead found this inflammatory response drove metastasis (83).

These results confirm *ZFP36* as an important component driving prostate cell identity and loss of *ZFP36* results in alternate transcriptional programs being activated and sustained. We and others have previously demonstrated the role that NF-κB plays in modulating response to androgen-deprivation therapy (3, 11, 84). The data presented here clearly demonstrate that loss of *ZFP36* increases resistance to anti-androgen therapy, at least in-part due to unregulated NF-κB activation. Further, our findings suggest that loss *of ZFP36* occurs in localized disease and is sufficient to induce multiple inflammatory pathways and phenotypic plasticity. This highlights the biological importance of *ZFP36* in PCa and provides the scientific basis to develop a new therapeutic strategy targeting NF-κB inhibitors to treat patients that present with PCa with low or absent *ZFP36* expression.

## Acknowledgements

We would like to thank Ms. Mariana Secundes for her technical assistance with mouse breeding and *in vivo* aging studies. We thank the support of the NKI Core Facility Molecular Pathology and Biobanking for human prostate tissue processing and analyses. This present study was supported by funding from the Department of Defense Idea Development Award (C.S.: W81XWH1910564); Department of Defense Early Career Investigator Award (K.M.: W81XWH1910305), National Cancer Institute (L.E.: R01CA207757, R01CA252468, R21CA257484), Oncode Institute (WZ) and the Dutch Cancer Society / Alpe d’HuZes (W.Z. and A.M.B: 10084).

## Author contributions

Conceptualization: LE, CJS, KLM, AAH Methodology: LE, CJS, KLM, AAH

Investigation: LE, KLM, AAH, BGF, JSN, SL, AMB, HvdD, IH, EMB, ST, DLB, SW, ATK, MK, JK, JTP, SY

Visualisation: KLM, AAH, SL, LE

Writing – original draft: LE, CJS, KLM, AAH

Writing – review and editing: LE, CJS, KLM, AAH, AGS, WZ, SL

## Supplementary Figure Legends

**Supplementary Figure 1: Genetics of prostate specific Pten/Ttp deleted GEMM mice.** (A) Schema of generated GEMM alleles including wild-type (WT), floxed (flox) and recombined (Δ) alleles. (B) Genotyping and recombination of PCR products from ear notch and prostate DNA samples of a PB-Cre4/Pten^f/f^/*Zfp36*^f/f^ mouse, highlighting PB-Cre4, Pten and *Zfp36* status.

**Supplementary Figure 2: Homozygous *Zfp36* loss results in progression to prostatic intraepithelial neoplasia.** (A) hematoxylin and eosin IHC and Krt18 (green) and alpha-SMA (red) IF staining of *Zfp36*+/- and *Zfp36*-/- murine dorsolateral prostate tissue 38 weeks. (B) Proliferation of PrEC-AR cells with and without *ZFP36*, +/- R1881 stimulation. (C) Proliferation and matching quantification of GEMM-derived 2D cell lines, n=3, *p<0.05, **p<0.005.

**Supplementary Figure 3: Validation of *in vivo* RNA-seq results are tumor cell autonomous.** (A) Principal component analysis plot of GEMM-derived 2D cell lines demonstrating separation of SKO (PBCre4:*Pten^f/f^)* and SKO-TTP (PBCre4:*Pten^f/f^*:*Zfp36^f/f^)* from PPKO (PBCre4:*Pten^f/f^*:*p53^f/f^)*, DKO (PBCre4:*Pten^f/f^*:*Rb1^f/f^)* and TKO (PBCre4:*Pten^f/f^*:*Rb1^f/f^*:*p53^f/f^)* samples. SKO-TTP samples cluster and separate from SKO control. (B) Gene set enrichment analysis of SKO-TTP versus SKO demonstrates significant enrichment for immune and inflammatory gene sets in the SKO-TTP group.

**Supplementary Figure 4: Loss of *Zfp36* drives castration resistance independent of *Tp53* and *Rb1* loss of function.** (A) *In vivo* tumor growth in GEMM-derived allograft models following surgical castration or sham castration at 7 days post-tumor implantation, n=5 mice per group, +/-1SD. (B) GEMM-derived 2D cell lines treated for 72 hrs with Nutlin-3 (5 µM) and Palbociclib (2 µM). GEMM-derived cell lines involving *Zfp36* loss generated for this manuscript were compared to previously published genetically defined murine models; *Pten^f/f^* (SKO), *Pten^f/f^*:*Rb1^f/f^* (DKO)*, and Pten^f/f^*:*Rb1^f/f^*:*p53^f/f^* (TKO) (48).

**Supplementary Figure 5: Functional effects of p65 inhibition in TTP-null GEMM-derived cell lines.** (A) Phosphorylated p65 protein expression in parental and modified cell lines. (B) Measurement of cell death in parental and modified cell lines treated with vehicle, DMAPT (5 µM), enzalutamide (5 µM) or the combination for 72 hours, +/- 1SEM. Inhibition of p65 restores enzalutamide sensitivity and reduces efficacy of DMAPT. (C) DMAPT dose-response cell death curves for parental and modified cell lines, with corresponding LD50 values (µM).

## Notes

### Competing Interest Statement

C. J. Sweeney is a stockholder in Leuchemix. All other authors declare no financial interests.

### Summary of Updates

Relevant data from two clinical trials has been added.

