## Supplementary figures and images for "Loss of tristetraprolin activates NF-κB induced phenotypic plasticity and primes transition to lethal prostate cancer"

### Supplemental Figures

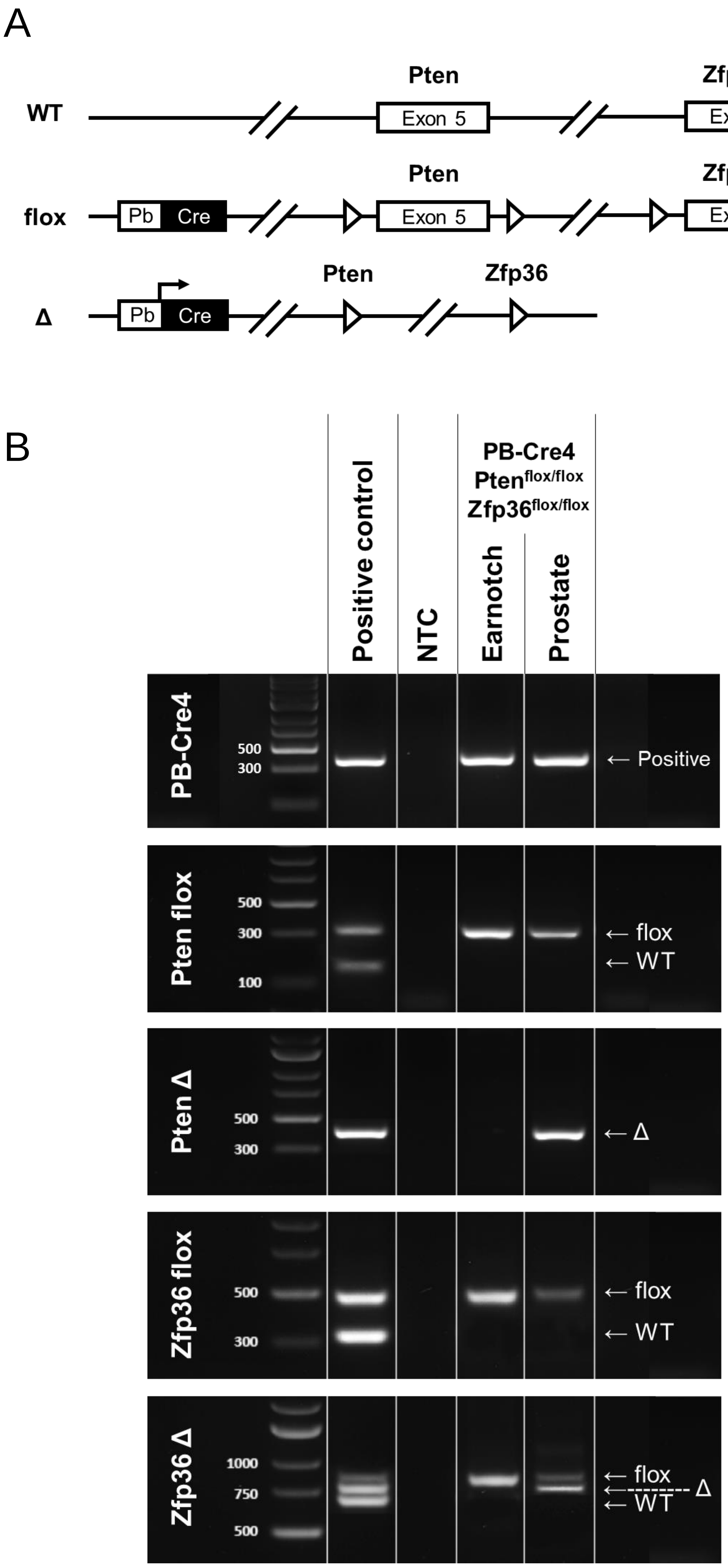

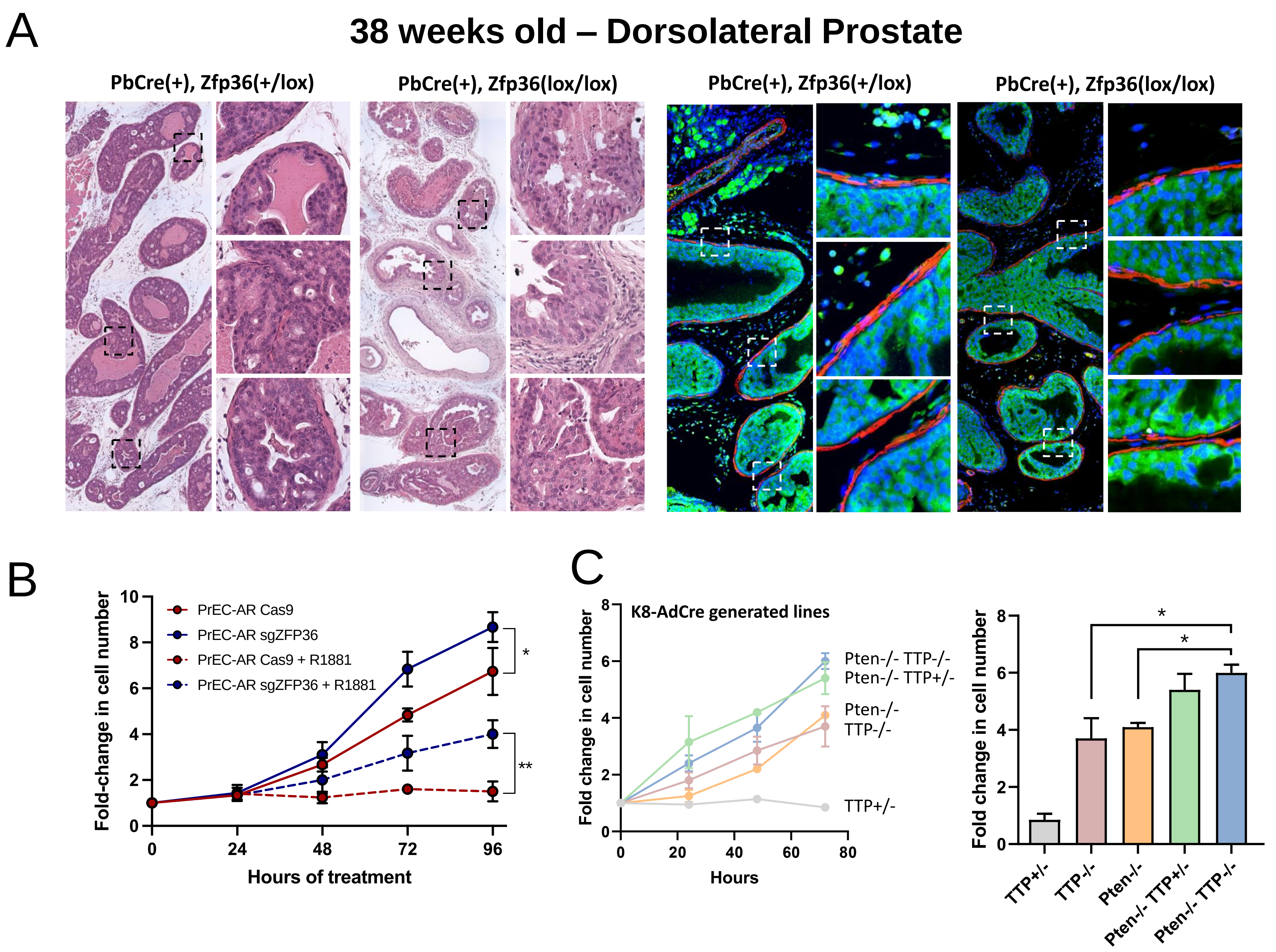

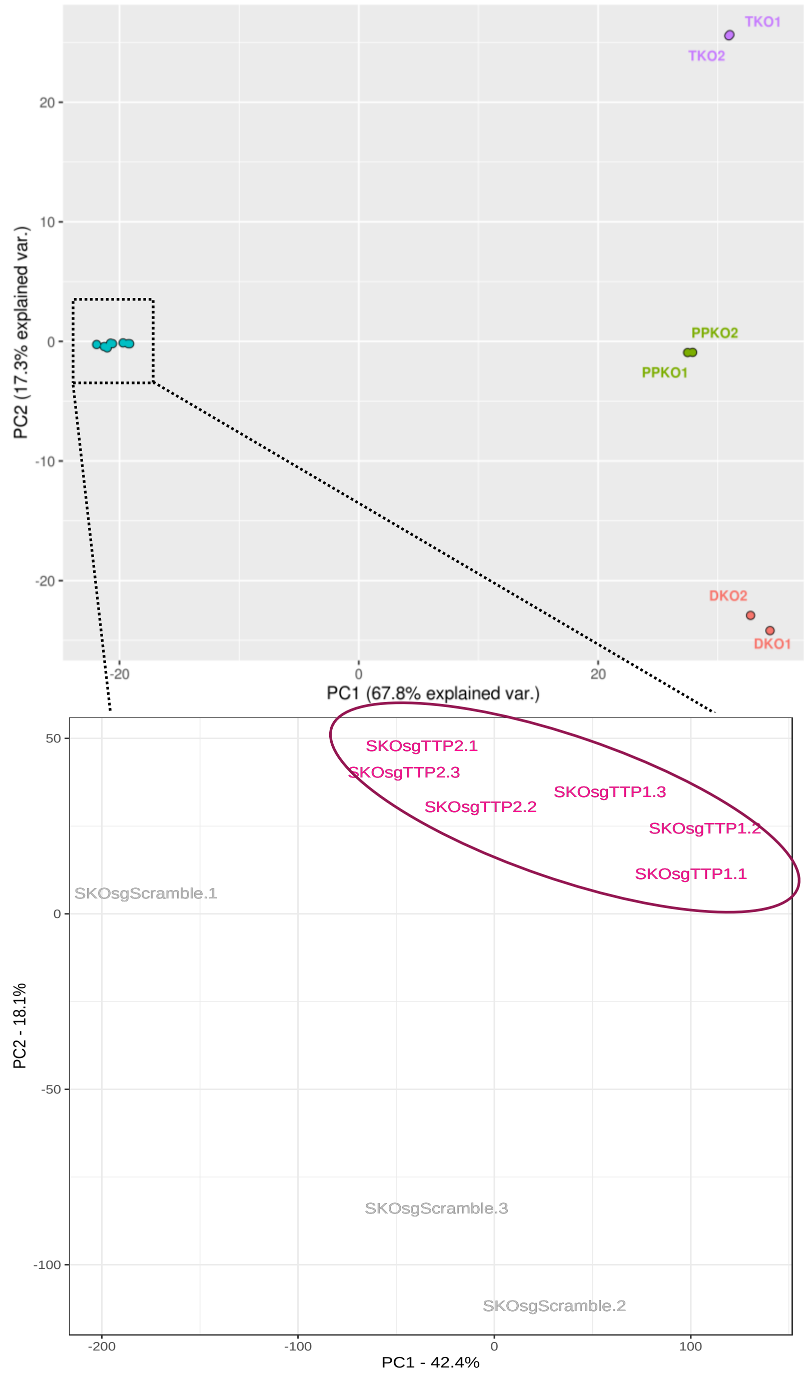

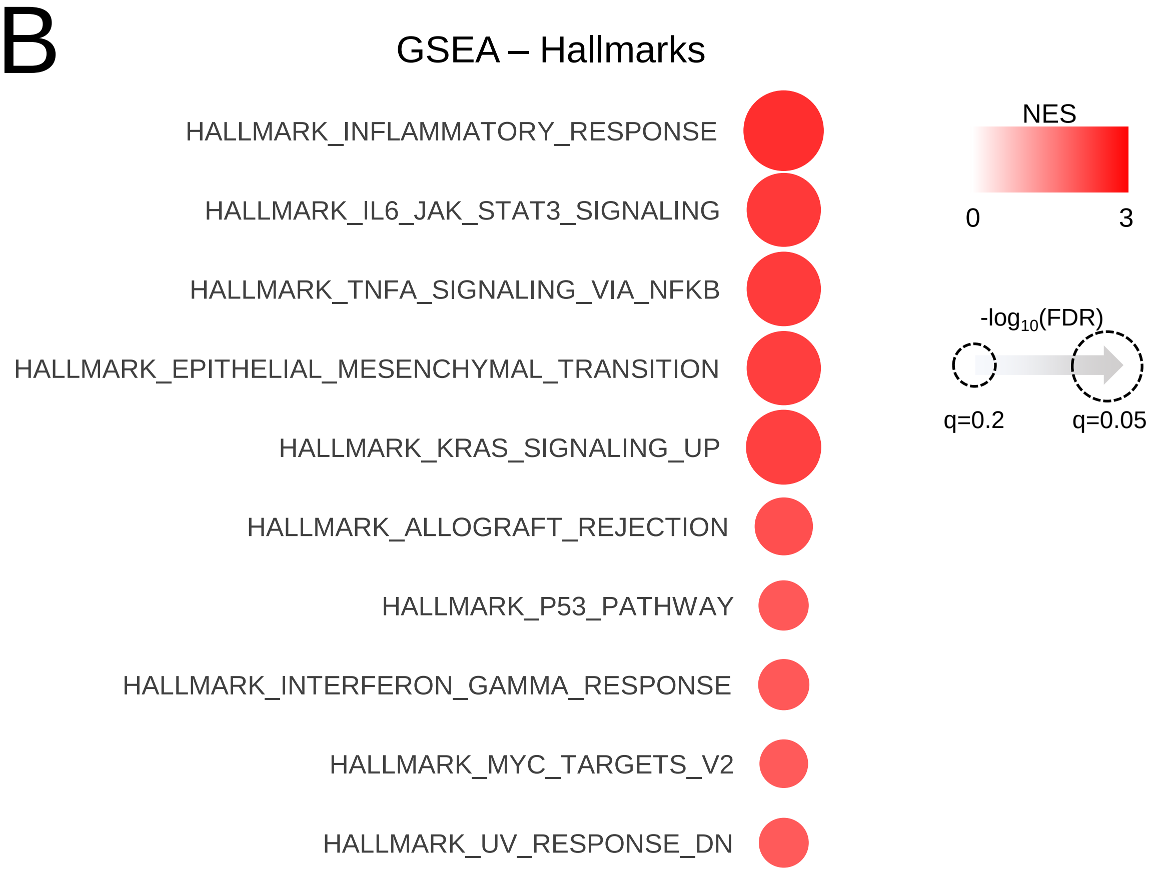

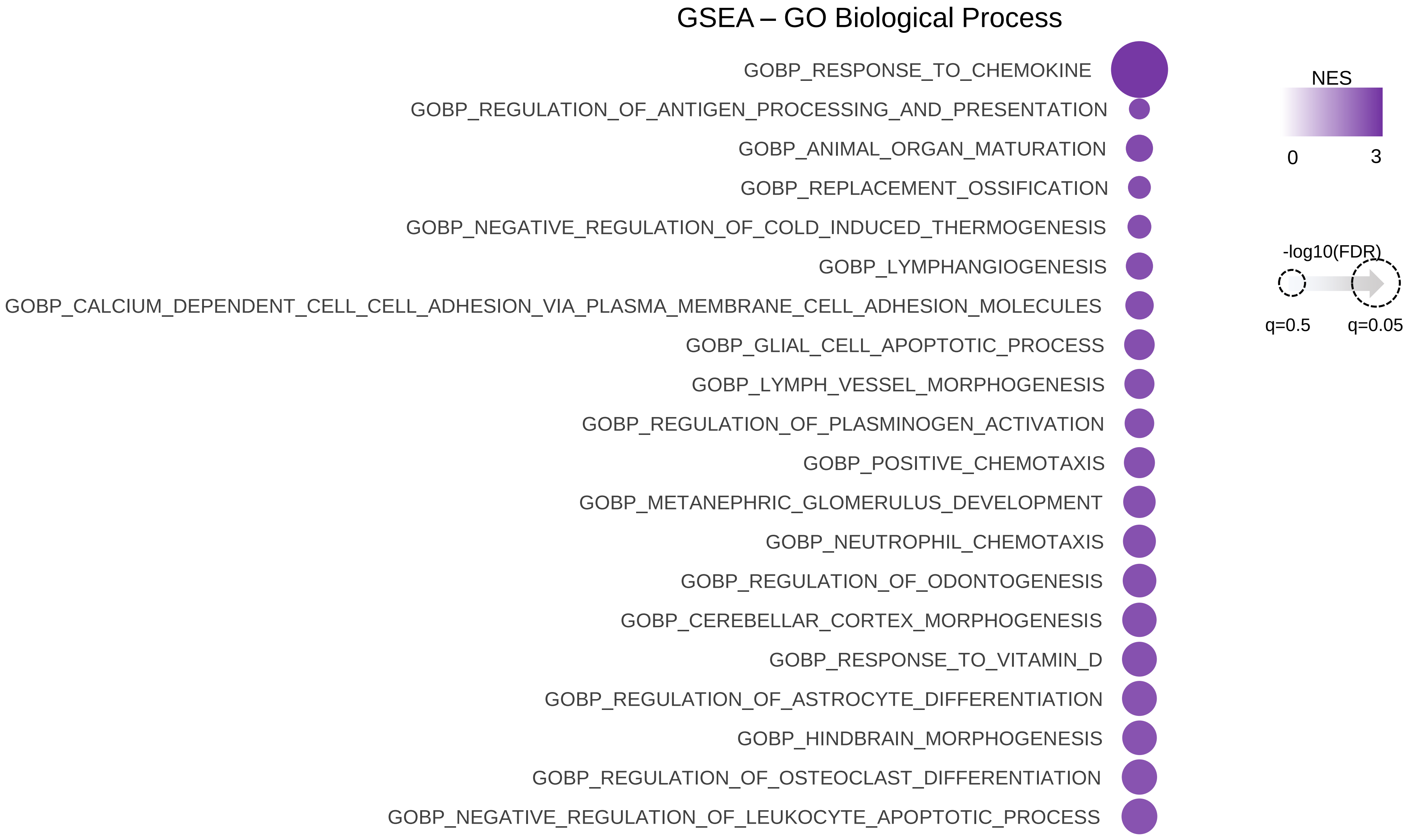

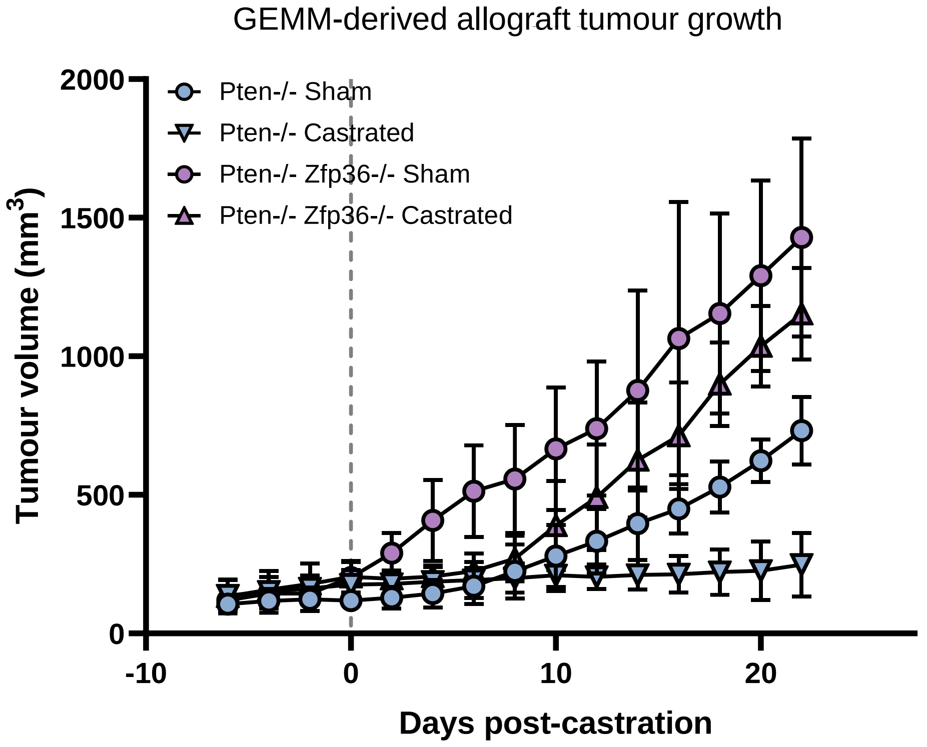

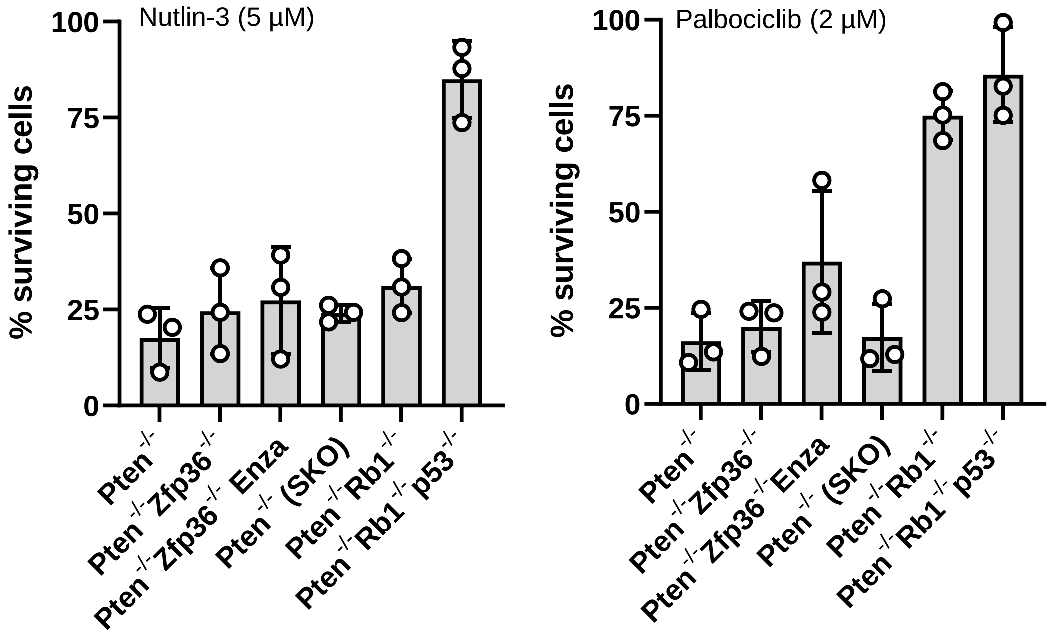

#### Figure S5

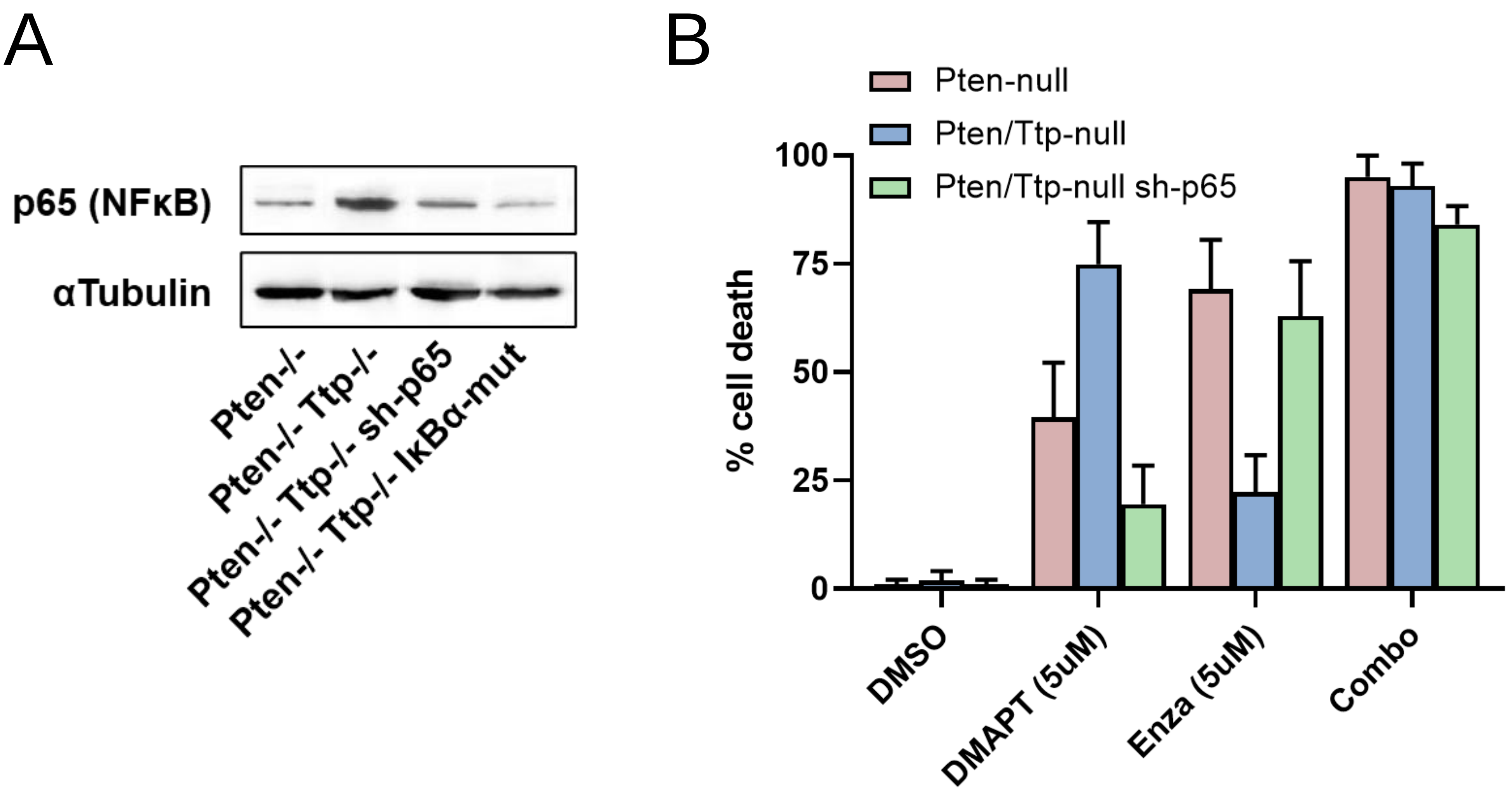


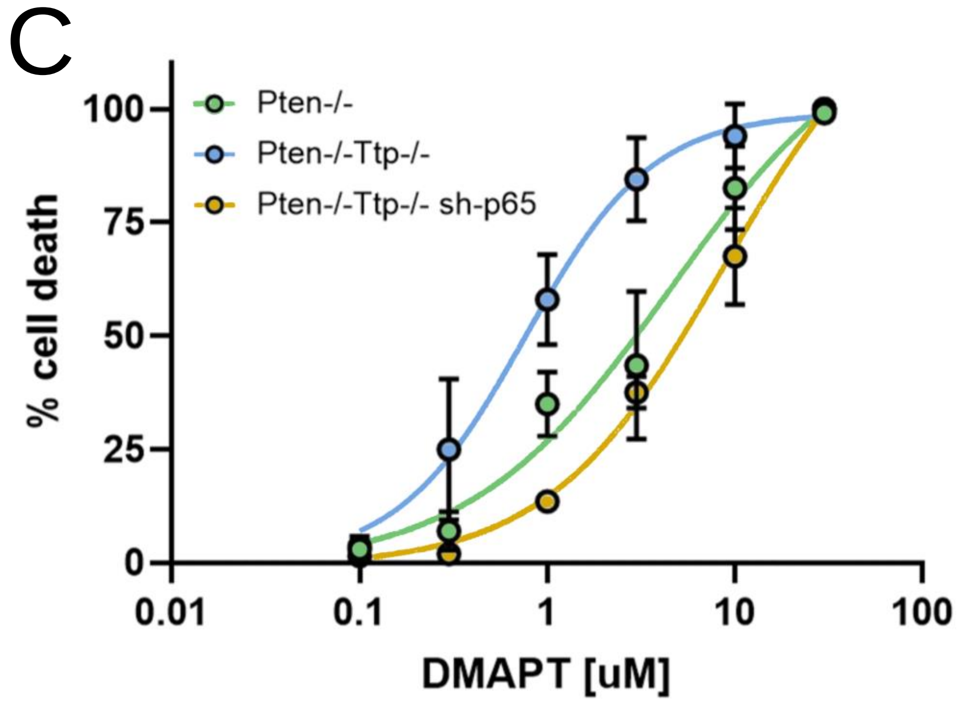


| Cell line | LD50 |
|---|---|
| Pten-/- | 4.52 |
| Pten-/- Ttp-/- | 0.76 |
| Pten-/- Ttp-/- sh-p65 | 8.66 |
